# Mitochondrial network structure controls cell-to-cell mtDNA variability generated by cell divisions

**DOI:** 10.1101/2022.06.27.497759

**Authors:** Robert C. Glastad, Iain G. Johnston

## Abstract

Mitochondria are highly dynamic organelles, containing vital populations of mitochondrial DNA (mtDNA) distributed throughout the cell. Mitochondria form diverse physical structures in different cells, from cell-wide reticulated networks to fragmented individual organelles. These physical structures are known to influence the genetic makeup of mtDNA populations between cell divisions, but their influence on the inheritance of mtDNA at divisions remains less understood. Here, we use statistical and computational models of mtDNA content inside and outside the reticulated network to quantify how mitochondrial network structure can control the variances of inherited mtDNA copy number and mutant load. We assess the use of moment-based approximations to describe heteroplasmy variance and identify several cases where such an approach has shortcomings. We show that biased inclusion of one mtDNA type in the network can substantially increase heteroplasmy variance (acting as a genetic bottleneck), and controlled distribution of network mass and mtDNA through the cell can conversely reduce heteroplasmy variance below a binomial inheritance picture. Network structure also allows the generation of heteroplasmy variance while controlling copy number inheritance to sub-binomial levels, reconciling several observations from the experimental literature. Overall, different network structures and mtDNA arrangements within them can control the variances of key variables to suit a palette of different inheritance priorities.

## Introduction

Mitochondria are vital bioenergetic organelles responsible for the majority of cellular energy production in eukaryotes [1, 2]. Due to their evolutionary history, mitochondria have retained small genomes [3–5] that encode genes central to their energy production [6, 7]. MtDNA in several taxa, including many animals, is subject to a high mutation rate relative to the nucleus, and mutations in mtDNA cause cellular dysfunction, and are involved in a range of human diseases [8, 9]. As mtDNA is predominantly uniparentally transmitted [10], the question arises of how eukaryotes avoid the gradual accumulation of mtDNA mutations, known as Muller’s ratchet [11].

The proportion of mutant mtDNA in a cell is usually referred to as heteroplasmy, and diseases are often manifest when heteroplasmy exceeds a certain level [12]. Eukaryotes may deploy a combination of strategies to generate cell-to-cell variability in inherited heteroplasmy [13–16], thus potentially generating offspring with heteroplasmies below the threshold [9, 17, 18]. For instance, in mammalian development, a developing female produces a set of oocytes for the next generation. Through an effective ‘genetic bottleneck’, oocytes with different heteroplasmies are generated [15]. This range of heteroplasmies means that some cells may inherit a lower heteroplasmy than the mother’s average. Across species, in concert with selection [19–24], this generation of variation allows shifts in heteroplasmy between generations [25, 26].

The mtDNA bottleneck has been suggested to incorporate a number of different mechanisms [27]. These include mtDNA depletion [28–30], and subpopulation replication [29, 31] in mammals, with a potential role for gene conversion in several other taxa [16], all coupled with random effects from partitioning of mtDNA at cell divisions. This partitioning is a focus of this report. Generally, when a cell divides, its mtDNA population will be partitioned between its daughter cells. Any deviation from precise deterministic partitioning (exactly half the mtDNA molecules of every genetic type go into each daughter) will likely lead to the daughter cells inheriting different heteroplasmy levels.

Models for this segregation have typically regarded the cellular mtDNA population as well-mixed, but the mitochondria containing the population have varied morphologies and dynamics in different cell types. Mitochondria can form cell-wide networks in some cell types, undergoing fission and fusion [32–36]. These dynamic structures, together with mtDNA turnover, are linked to quality control and genetic dynamics of mtDNA populations [24, 37, 38]. In both somatic tissues [39–42], or the germline [16, 23], the balance between fusion, fission and selective degradation of individual dysfunctional mitochondria has emerged as an important influence on mtDNA populations [39, 41, 43–45].

The important effects of heterogeneous spatial distributions on noise in other cell biological contexts are being increasingly recognised [46]. The central importance of mitochondria in cell metabolism and bioenergetics mean that variability in their physical and genetic inheritance can influence many other, also noisy, downstream processes [47, 48]. Quantitative progress modelling the spatial influence of these mitochondrial dynamics on mtDNA quality control is advancing [39–42, 45]. In particular, the role of network structure in generating cell-to-cell mtDNA variability via modulating mtDNA turnover has been addressed with recent stochastic modelling [16, 49]. These studies report that cell-to-cell mtDNA variability increases with the proportion of fragmented mitochondria in the cell, the rate of turnover, and the length of the cell cycle. Moreover, turnover itself may increase due to a highly fragmented network, with a highly fused network effectively masking mitochondria marked for degradation. However, the influence of mitochondrial network structure on the inheritance of mtDNA during cell divisions remains less studied. Although the mitochondrial network may fragment prior to fragmentation, allowing individual mitochondria to diffuse, or actively mix [50], the network structure prior to division may exert substantial influence on the distribution of mtDNAs in the parent cell [32], ultimately reflecting in daughter cell statistics. Partitioning at cell divisions even in the absence of spatial substructure can constitute an important source of cell-to-cell variability [51, 52]. In yeast, mtDNA inheritance occurs with finer-than-random (binomial) control over the number of mitochondrial nucleoids [53], suggesting that physical mechanisms must exist to exert this control. As mtDNA resides in nucleoids that are physically distributed – potentially heterogeneously – throughout the mitochondria of the cell [54], we set out to explore how different physical structures of mitochondria may shape mtDNA inheritance.

## Results

### Network inclusion with genetic bias increases cell-to-cell variability

To build intuition about the influence of network structure on mtDNA inheritance, we first considered a simple computational model for the spatial distribution of mitochondria and mtDNA within a cell (see Methods). This model consists of a random network structure, with a tunably heterogeneous distribution, simulated within a circular cell (the dimensionality of the model does not affect the statistical considerations of partitioning) (see Fig. 1). The model network structure is not intended to perfectly match the details of real mitochondrial networks, but rather as a general framework to understand spatial substructure in the cell. The mother cell has *N*_0_ mtDNAs, a proportion *h* of which are mutants (*h* is heteroplasmy).

**Figure 1:**
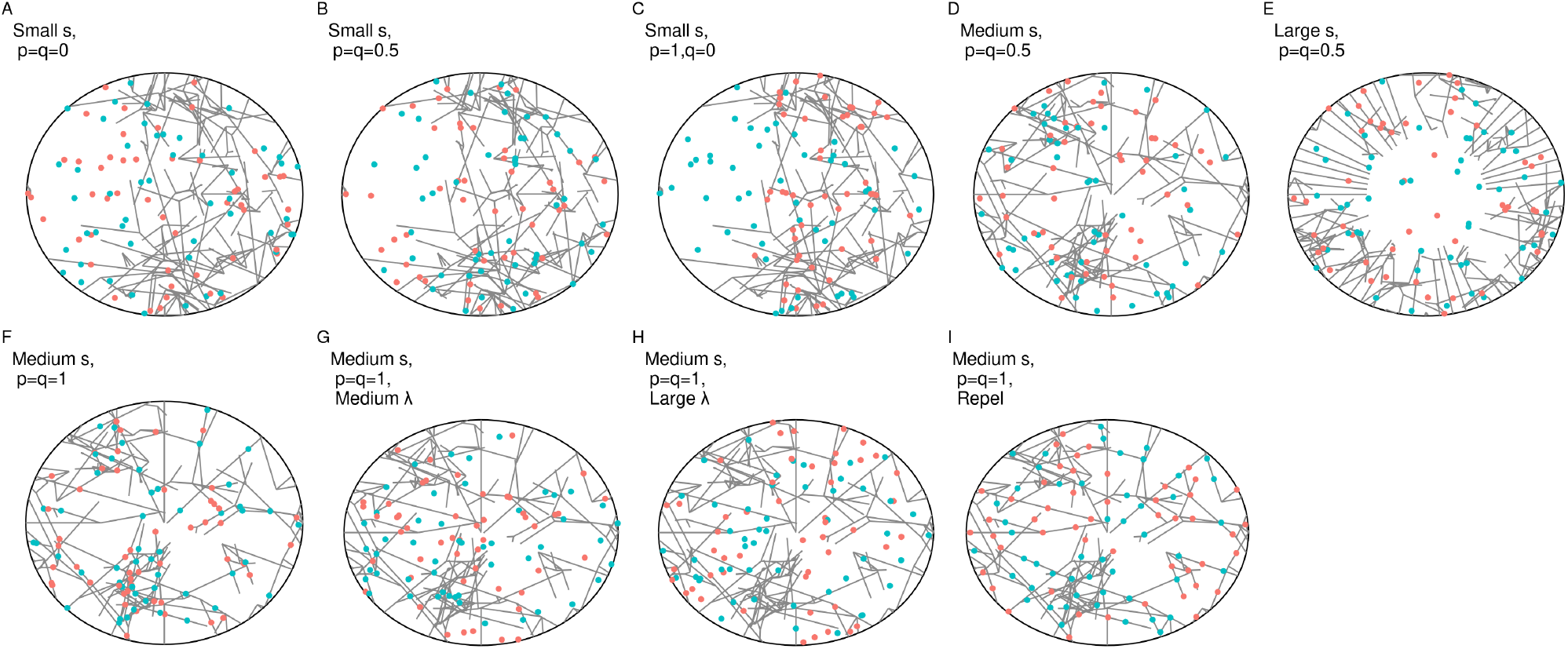
Snapshots of computationally generated networks and mtDNA arrangements with tunable physical and genetic parameters. Networks were generated by a elongation and branching process initialized from a number of seed points (s) uniformly distributed on the perimeter of the cell. Small seed numbers *s* (A-C) usually resulted in heterogeneous network structures, with marked differences in network density across the cell; for increasing seed numbers (D-F), networks were more uniformly distributed throughout the cell. Wild type (WT; red) and mutant type (MT; blue) mtDNA molecules were distributed into networks according to the proportions *p* (WT) and *q* (MT). In different model variants, diffusion with scale parameter λ was applied to mtDNAs prior to division to model network fragmentation and subsequent motion (G-H; original network shown for reference), and mtDNAs were placed with a repulsive interaction inducing greater-than-random spacing (I).

To reflect the fact that different mtDNA types may have different propensities to be included in the reticulated network [43], we use *p* and *q* to describe the proportions of wildtype and mutant mtDNAs respectively that are contained in the network; the remainder are in fragments in the cytoplasm (Fig. 1). Hence, *p* > *q* means that wildtype mtDNAs are more likely to be contained in the network and mutant mtDNAs are less likely to be included; *p* = *q* means that both types are equally likely to be in a networked state. Network placement can follow various rules (described below) and mtDNA positions may be subsequently perturbed, but we begin with random placement in the network and no subsequent motion prior to division. We then divide the model cell and explore the statistics of mtDNA copy number and heteroplasmy in the daughter cells. We first consider symmetric partitioning, so that the initial cell is physically halved to produce two daughters; we relax this assumption and consider asymmetric partitioning later.

To explore the influence of mtDNA network placement in the parent cell on the mtDNA statistics of the daughter, we varied network inclusion probabilities *p* and *q* (Fig. 1A-C) and network heterogeneities (Fig. 1C-E) and observed copy number and heteroplasmy variance in daughter cells after division (Fig. 2). We found a clear pattern that *V*(*h*) takes minimum values when *p* = *q*, that is when network inclusion probabilities are equal for mutant and wildtype mtDNA. When the two differ, *V*(*h*) increases, with highest values occurring when the majority mtDNA type is exclusively contained in the network and the minority type exclusively in the cytoplasm.

**Figure 2:**
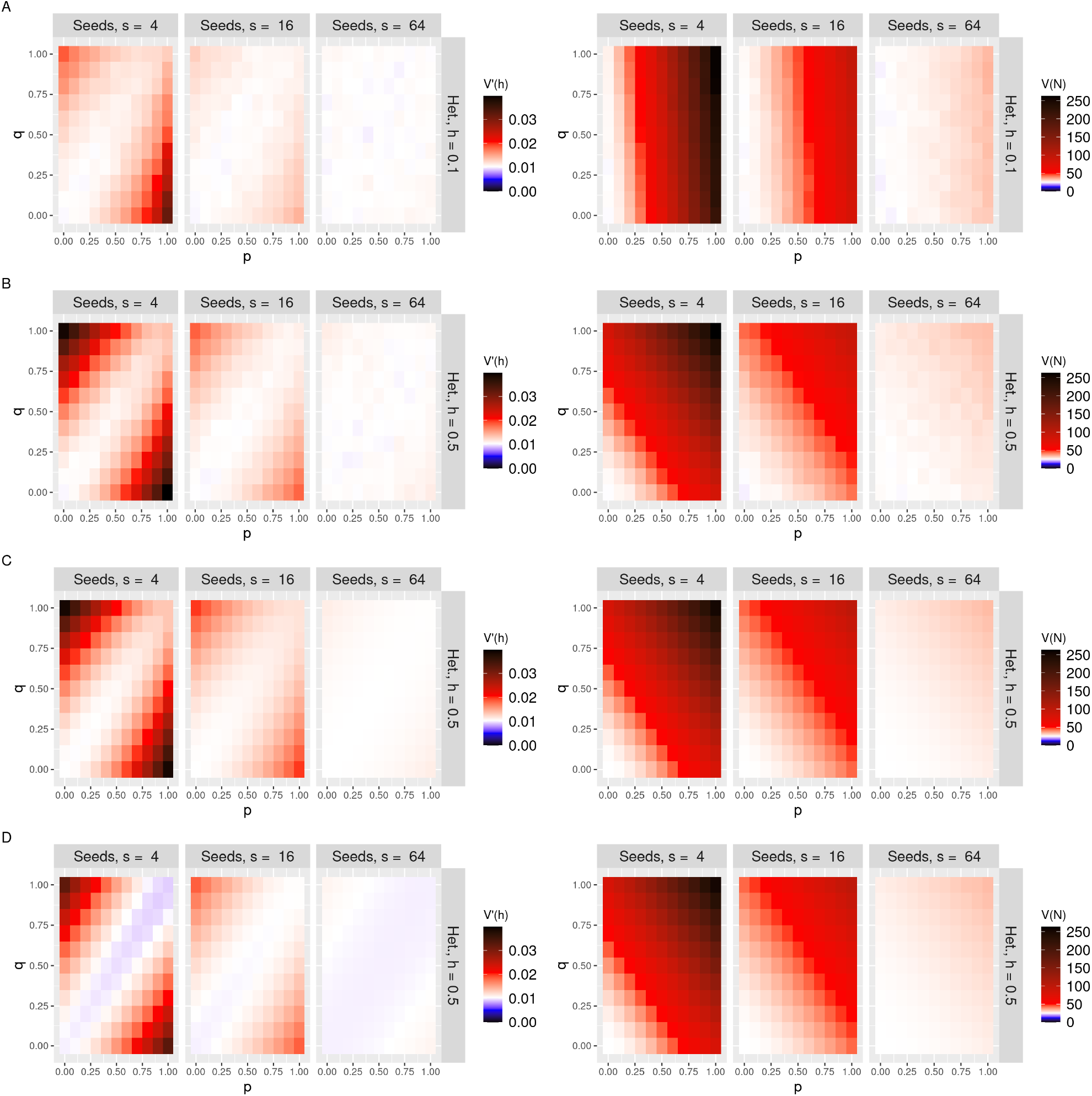
Genetic bias and physical heterogeneity in mitochondrial networks induce cell-to-cell mtDNA variability in symmetric cell divisions. Normalised heteroplasmy variance *V′*(*h*) (left column) and copy number variance *V*(*N*) (right column) for mtDNA randomly distributed in networks. **A**: simulations for *h* = 0.1; **B**: simulations for *h* = 0.5; **C**: sum over state variables for *h* = 0.5; **D**: first-order Taylor expansion for *h* = 0.5. The three columns to a facet give decreasing network heterogeneity, expressed via different seed numbers, 4, 16 and 64 (more seed points give a more homogeneous network). In each panel, variances are given for different values of wild and mutant type network inclusion parameters *p* (horizontal axis) and *q* (vertical axis). White baseline reflects the null case from the analytic sum without any network inclusion.

This result may seem counterintuitive at first glance: when the network is highly heterogeneous, it might be expected that including all mtDNAs there would maximise variance. This is true for copy number variance (Fig. 2), but not for heteroplasmy variance, because including both types equally induces correlation in their inheritance and lowers variance. The maximum *V*(*h*) is achieved by embedding the majority type in the high-variance network and having the minority type in the uncorrelated cytoplasm.

### Statistical models of mtDNA inheritance capture the variance induced by cell divisions

To further explore this behaviour, we constructed a statistical model for this inheritance process (see Methods). Briefly, we consider four state variables describing the mtDNA population of a daughter cell after a mother divides: *W_n_*,*W_c_*,*M_n_*,*M_c_*, for the wildtype (*W*) and mutant (*M*) mtDNAs contained in a reticulated mitochondrial network (_*n*_) or in fragmented mitochondrial elements in the cytoplasm (_*c*_). An additional variable *U* describes the proportion of network mass inherited by the daughter cell. Given a particular value *u* for this proportion, the mtDNA profile inherited by the daughter follows

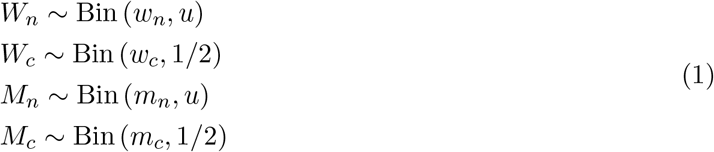

where *w_n_* = *p*(1 – *h*)*N*_0_,*w_c_* = (1 – *p*)(1 – *h*)*N*_0_, *m_n_* = *qhN*_0_, *m_c_* = (1 – *q*)*hN*_0_. We model the proportion *u* of network mass inherited by a daughter with a beta-distributed variable *U*, with variance *V*(*U*) allowed to vary to describe different partitioning regimes of network mass. To compare to simulations, we fit the beta distribution parameters to match the simulated inherited network variance. As *W_n_* and *M_n_* are then drawn from the compound distribution that is binomial with a beta-distributed probability, they follow beta-binomial distributions. We are interested in the inherited copy number *N* = *W_n_* + *W_c_* + *M_n_* + *M_c_* and heteroplasmy *h* = (*M_n_* + *M_c_*)/*N*.

We can numerically extract moments for the system through summing over state variables and calculating expectations, for example,

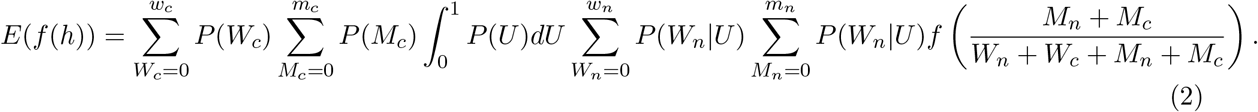

Figs. 2 and 3 demonstrates good correspondence between simulations and statistics using Eq. 2 for copy number and heteroplasmy variance (*V*(*X*) = *E*(*X*^2^) – *E*(*X*)^2^). However, as these large sums do not admit much intuitive analysis, we sought other approaches to learn the forms of pertinent statistics of the inherited mtDNA population.

**Figure 3:**
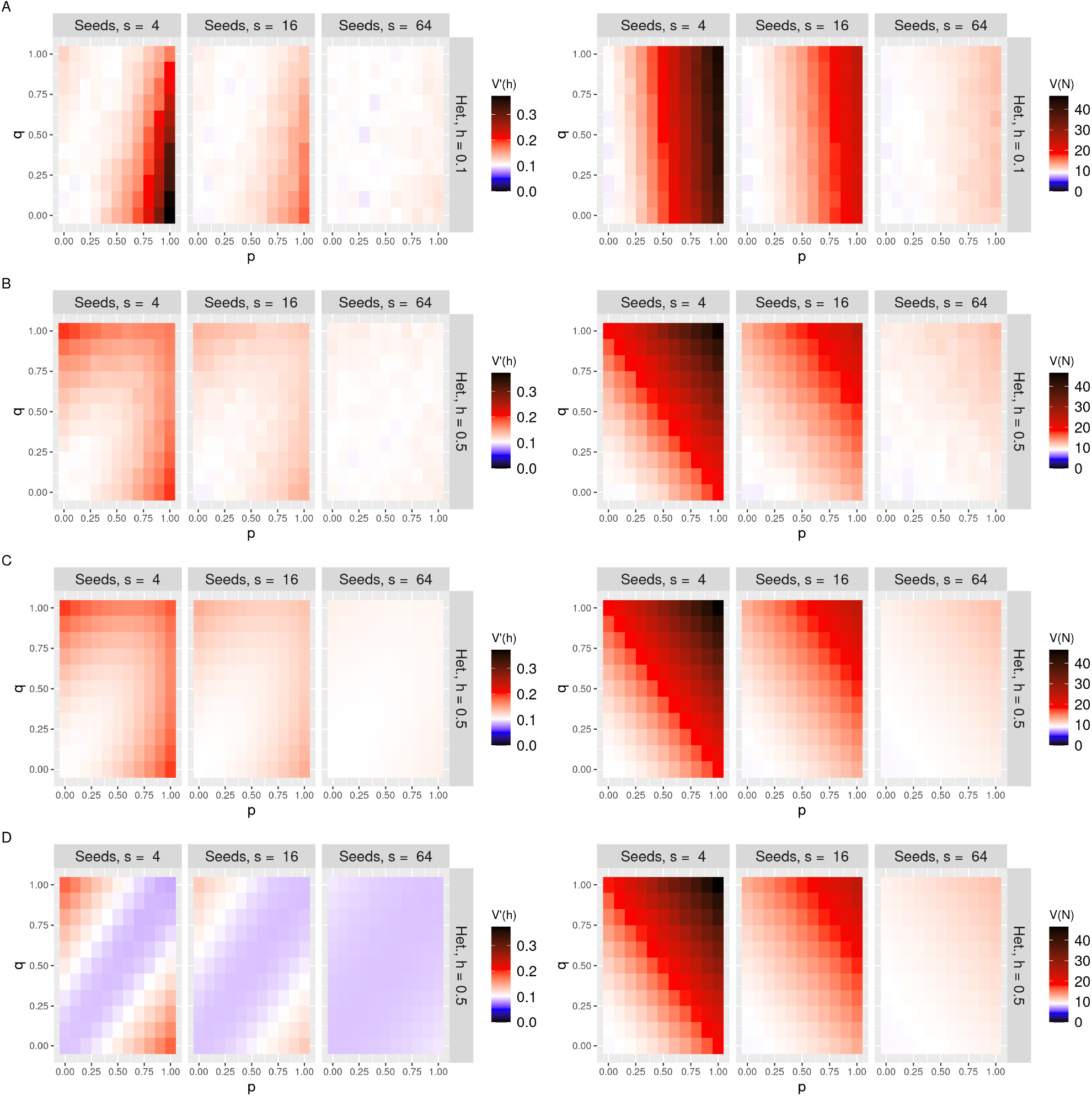
Asymmetric cell divisions can induce large cell-to-cell variability in mtDNA quality. Normalised heteroplasmy variance *V′*(*h*) (left column) and copy number variance *V*(*N*) (right column) for mtDNA randomly distributed in networks. The daughter of interest inherits 10% of the parent cytoplasm. **A**: simulations for *h* = 0.1; **B**: simulations for *h* = 0.5; **C**: sum over state variables; **D**: first-order Taylor expansion). The three columns to a facet give decreasing network heterogeneity, expressed via different seed numbers, 4, 16 and 64 (more seed points give a more homogeneous network). In each panel, variances are given for different values of wild and mutant type network inclusion parameters *p* (horizontal axis) and *q* (vertical axis). White baseline reflects the null case from the analytic sum without any network inclusion.

The mean and variance of inherited number *N* are readily derived using the laws of iterated expectation and total variance to account for the compound distribution of networked mtDNA (Appendix):

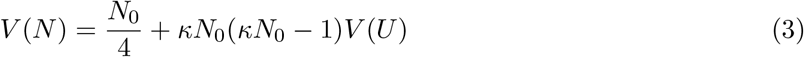

where *κ* = *p*(1 – *h*) + *qh* denotes the proportion of total mtDNA placed in the network. Hence *V*(*N*) experiences an extra, *V*(*U*)-dependent term in addition to the expected *N*_0_/4 result for a purely binomial distribution. This term is quadratic in the proportion *κ* of mtDNA in the network. This expression well captures the results from simulation and Eq. 2 (Fig. 2).

For heteroplasmy variance, as the ratio of random variables, we cannot extract an exact solution and must instead use a Taylor-expanded approximation (see Methods) to obtain

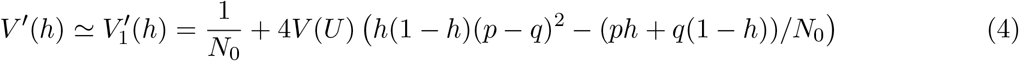

Eq. 4 allows some informative analysis. First, we qualitatively see that an additional, *V*(*U*)-dependent term is introduced compared to the binomial case (which gives 1/*N*_0_), illustrating the influence of network heterogeneity on heteroplasmy variance. For large *N*_0_, this network term is dominated by the first term in brackets in Eq. 4, which is quadratic in (*p* – *q*), the difference in inclusion probabilities for the different types of mtDNA. For *p* ≠ *q*, the network is genetically biased towards one of the types, to which there is associated an increase in *V′*(*h*). For *p* = *q*, the network is unbiased, with associated prediction 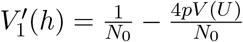 – that is, a small negative change from the binomial case. However, this negative shift is in fact an artefact of the imperfect Taylor approximation we use (see below and Appendix), and the *p* = *q* case in fact resembles the binomial case with a slight increase (captured by a higher-order approximation, see Appendix) at higher *p* (Fig. 2).

The more useful prediction under this approximate model concerns the maximum normalised heteroplasmy variance achievable – when the majority mtDNA type is completely contained in the network and the minority type completely excluded from it – is given for example by setting *p* = 1, *q* = 0 for *h* ≤ 0.5:

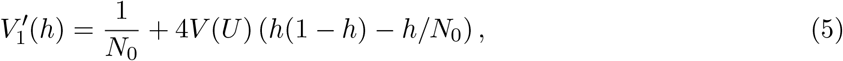

with a *V*(*U*)-dependent term scaled by *h*(1 – *h*) (in most cases the *h/N*_0_ term will be negligible), illustrating that genetic bias in network inclusion can substantially increase heteroplasmy variance in proportion to inherited network variability.

This picture does not completely capture the simulation and exact results, where we see a small increase in (normalized) heteroplasmy variance for non-biased increases in inclusion probabilities, instead of a decrease. This reflects the approximate nature of the Taylor expansion process used to derive Eq. 4; in the Appendix we show that a second-order expansion provides a compensatory diagonal term (Fig. 8), but in general we will need further terms in the expansion to perfectly match the true behaviour. In the Appendix we further show that all higher-order moments and covariances of the mtDNA copy numbers are well captured by theory; it is their combination into an estimate for the moments of a ratio (heteroplasmy) that is challenged here. We will observe below that more instances of this system also challenge this Taylor expansion approach, which has been employed previously [27, 55, 56] – generally, accounting for physical structure induces correlations between mtDNA types that are hard to capture with the Taylor expression (see Discussion).

**Figure 4:**
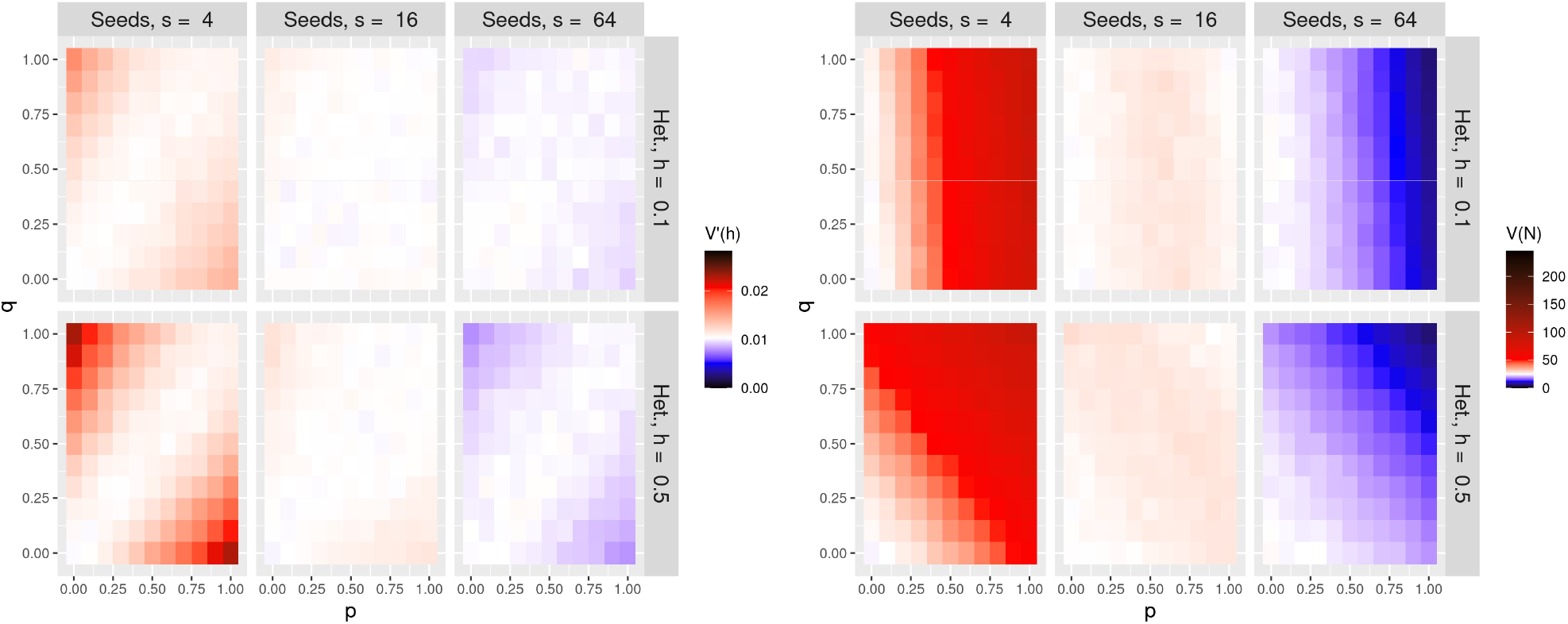
Repulsion between mtDNAs in the network can increase and decrease cell-to-cell variability from cell divisions. Simulated normalised heteroplasmy variance *V′*(*h*) (left column) and copy number variance *V*(*N*) (right column) for mtDNAs with mutual repulsion within the network, under symmetric cell divisions. Rows show different values of initial mutant proportion, with *h* = 0.1 in the top row and *h* = 0.5 in the bottom row. The three columns to a facet give decreasing network heterogeneity, expressed via different seed numbers, 4, 16 and 64 (more seed points give a more homogeneous network). In each panel, variances are given for different values of wild and mutant type network inclusion parameters *p* (horizontal axis) and *q* (vertical axis). White baseline reflects the null case from the analytic sum without any network inclusion.

**Figure 5:**
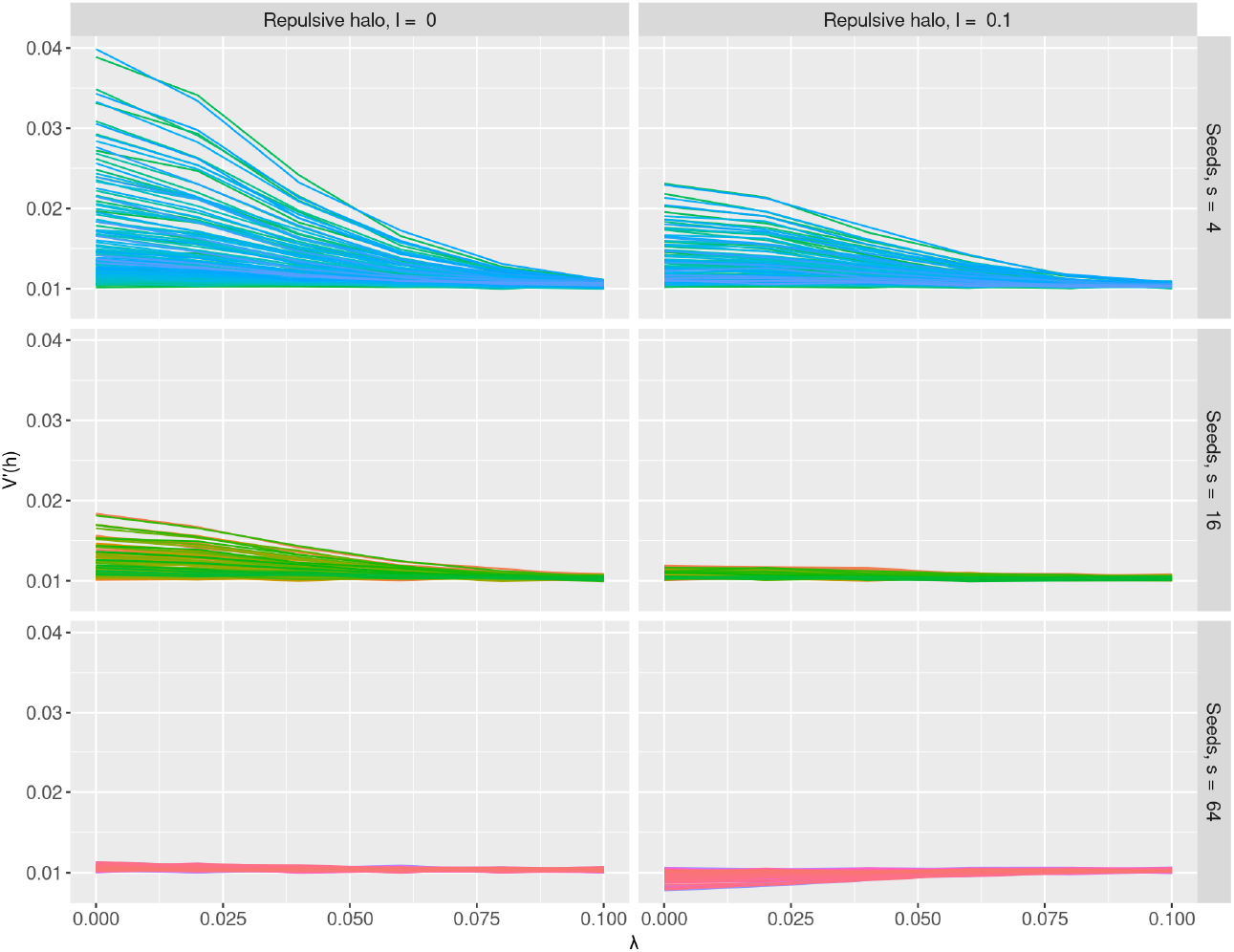
Network fragmentation and diffusion reverse the variance induced by mitochondrial networks. Normalised heteroplasmy variance *V′*(*h*) after repeatedly perturbing each mtDNA molecule with mean displacement λ from their original positions, resulting in a total mean displacement of around 10λ. The left column shows *V′*(*h*) for random mtDNA placement in the network; the right column shows *V′*(*h*) for repulsive mtDNA placement in the network. The three rows show decreasing levels of network heterogeneity, expressed via different seeds numbers (more seed points give a more homogeneous network). As the diffusion strength increases, effects on cell statistics due to the network is washed away regardless of the underlying mtDNA distribution model.

**Figure 6:**
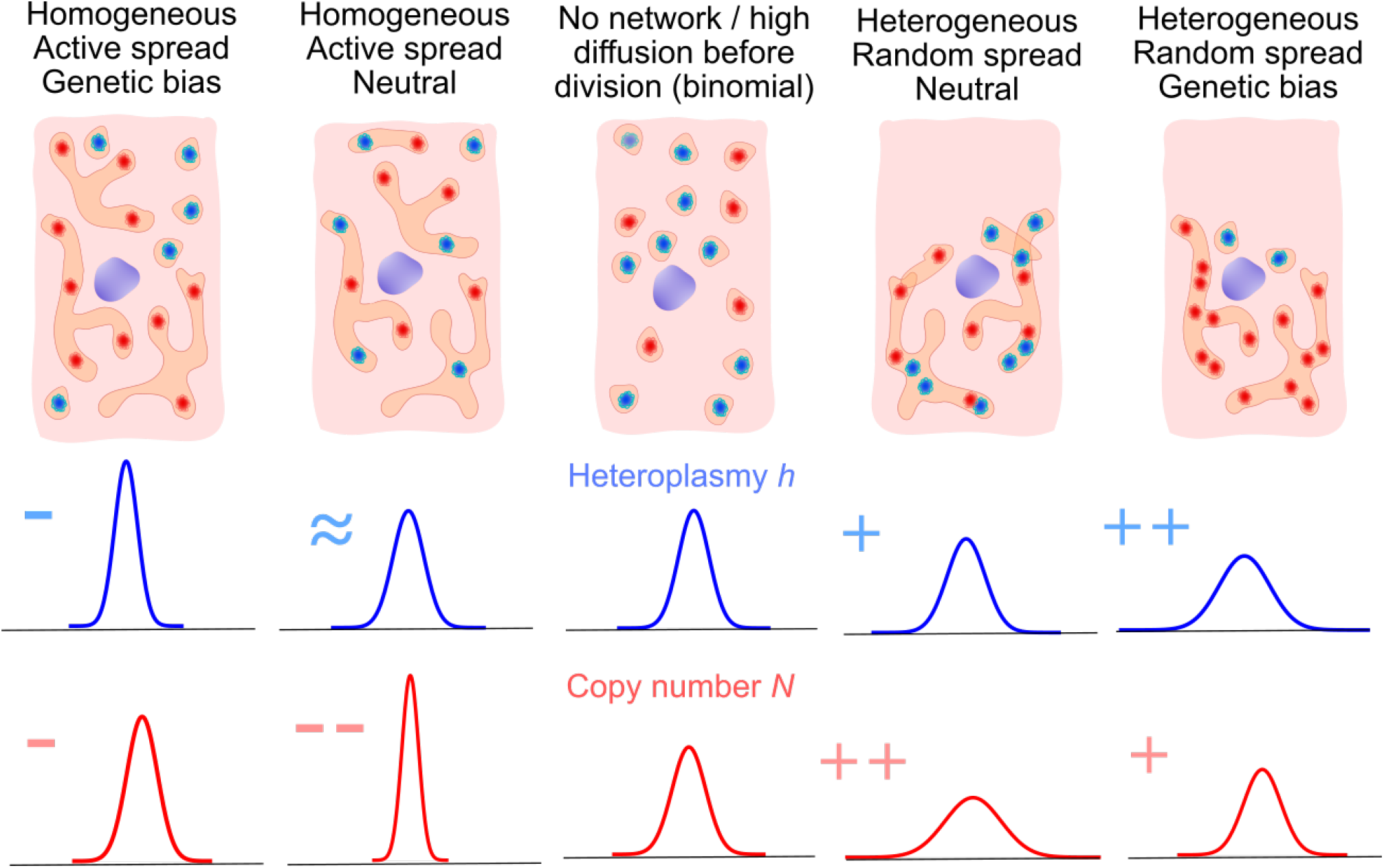
Illustration of the range of influences of network structure and contents on mtDNA statistics from partitioning. MtDNA molecules distributed randomly through the cytoplasm are segregated binomially (centre); the influence of network structure (heterogeneous or homogeneous distributions) and mtDNA inclusion (genetically neutral or biased) changes cell-to-cell variability in different ways: Depending upon genetic bias and network heterogeneity, networks can both increase and cell-to-cell mtDNA variability in copy number and heteroplasmy.

**Figure 7:**
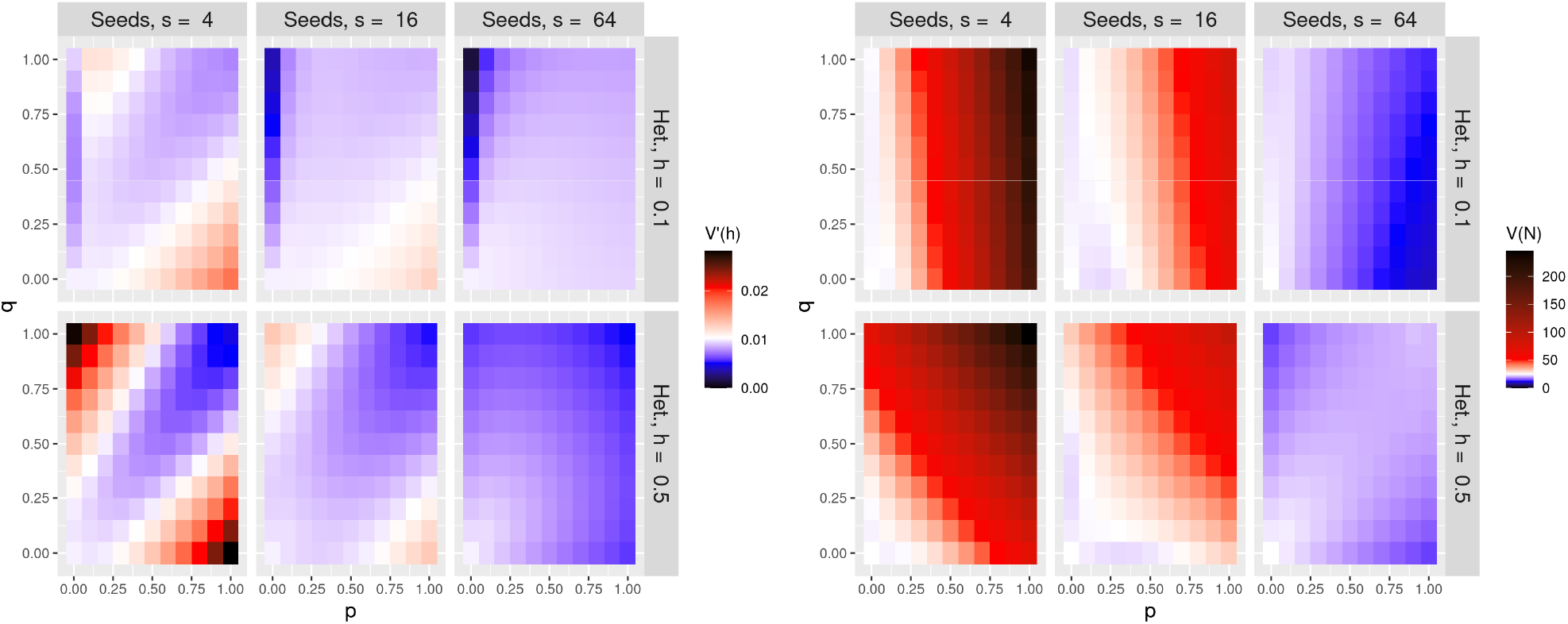
Approximate model predictions for repulsive interactions between mtDNAs in the network. *V′*(*h*) (left column) and *V*(*N*) (right column) under an approximate model for repulsive mtDNA placement in the network. Qualitatively trends in behaviour are captured, but the magnitudes of the effects involved differ from simulated results (see text).

**Figure 8:**
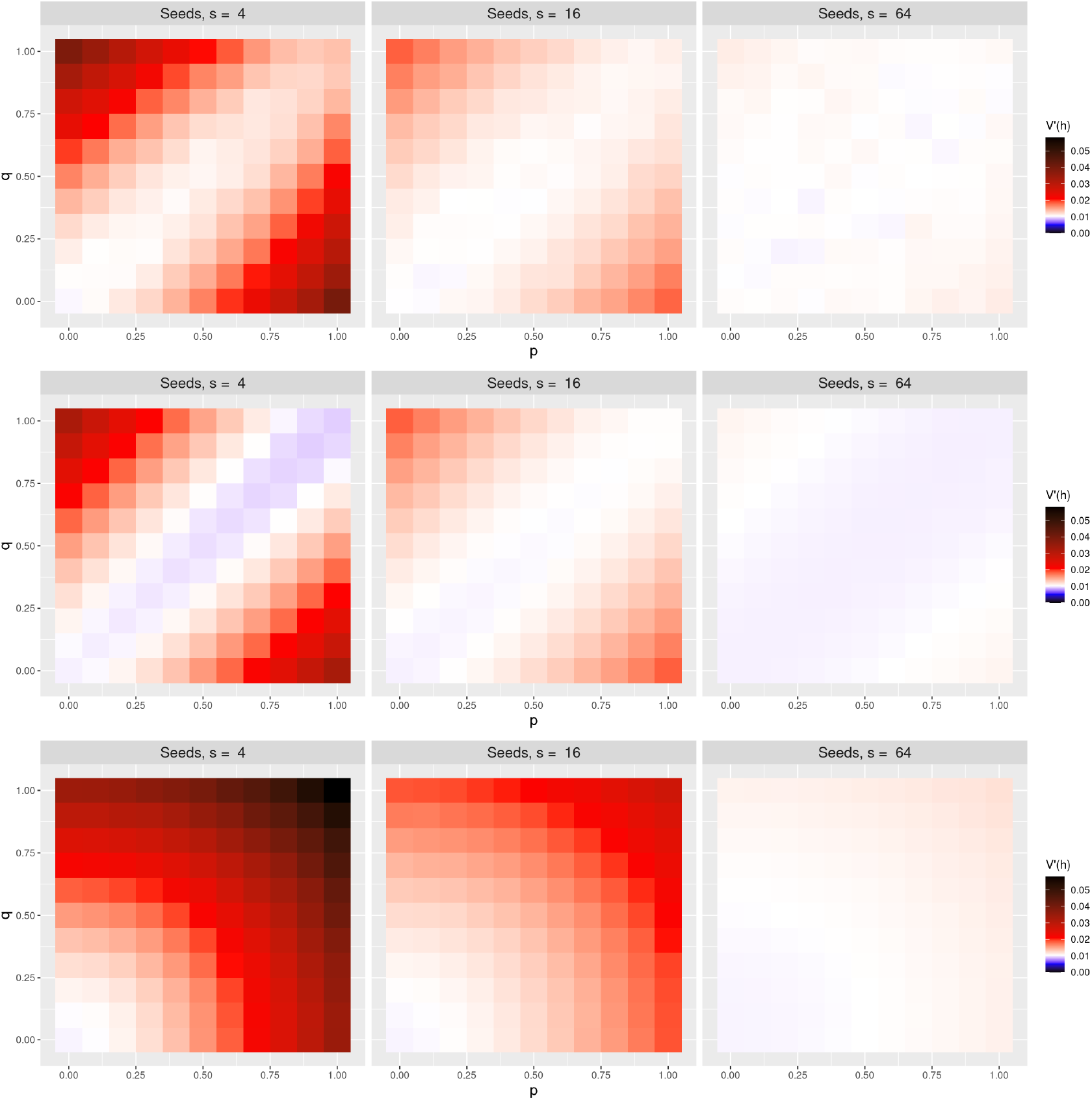
Comparison of simulation results with analytic model results for first and second order Taylor expansion. By row, simulation, first, and second order analytic results for *V′*(*h*) for random placement of mtDNAs in the network. The first order theory in the second row produces results similar to our simulation results on the off-diagonal, but fails to reproduce the increase observed along the diagonal. The second order theory, while loosely retaining the same structure on the off-diagonal as the first order theory, overestimates this increase along the diagonal. We expect that our model would captures this behavior if we were to derive even higher order terms.

### Asymmetric cell divisions induce more mtDNA variability

The above model has assumed that the cell divides symmetrically, with half the cell volume inherited by each daughter. To generalise to asymmetric cell divisions, we next asked how the proportion of inherited cell volume *p_c_* (*p_c_* = 1/2 in the symmetric case) influences the mtDNA statistics in daughters. Clearly, the expected copy number will differ if the inherited proportion differs. The expressions above generalise to

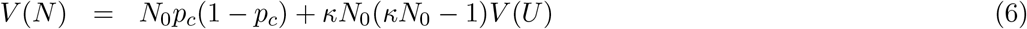

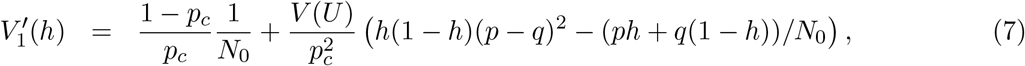

with the result that asymmetric cell divisions can generate large increases in cell-to-cell variability for the smaller daughters (Fig. 3). Small number effects are at play here, with a smaller sampling of the initial cell inevitably leading to greater relative variance. A decrease in *p_c_* from the symmetric case *p_c_* = 1/2 to *p_c_* = 0.1 results in a near order-of-magnitude increase in the maximum normalised heteroplasmy variance. Asymmetric divisions also further challenge the Taylor expansion approach, with a systematic underestimation of heteroplasmy variance apparent using this approximation (copy number variance remains well captured).

### MtDNA self-avoidance tightens mtDNA copy number control and can reduce heteroplasmy variance

We next asked whether the variances of copy number *V*(*N*) and heteroplasmy *V*(*h*) could be reduced below their ‘null’ binomial value through cellular control. To this end, we modelled self-avoidance of mtDNA molecules within the network (Fig. 1I), reasoning that such controlled arrangement may allow a more even spread of mtDNAs within the network, and correspondingly lower variability. To accomplish this self-avoidance within our model, we enforce a ‘halo’ of exclusion around each mtDNA placed within the network, so that another networked mtDNA cannot be placed within a distance *l* of an existing one.

Copy number variance *V*(*N*) is decreased substantially by self-avoidance (Fig. 4). In the case of a homogenous network and high proportions of mtDNA network inclusion, this decrease can readily extend below the binomial null model, allowing more faithful than binomial inheritance, as reported in yeast [53]. This sub-binomial inheritance requires both an even network distribution and mtDNA self-avoidance, and hence two levels of active cellular control. Under these circumstances, mtDNA molecules are evenly spread throughout the cell volume, and their inheritance approaches a deterministic proportion of the inherited volume fraction *p_c_*.

The effect of mtDNA self-avoidance on heteroplasmy variance is more complicated. For highly heterogeneous network distributions, heteroplasmy variance follows the same qualitative pattern as for the non-repulsive case, with higher variances achieved when network inclusion discriminates wildtype and mutant types. However, for homogeneous network distributions, the opposite case becomes true. Here, network inclusion discrimination *lowers* the heteroplasmy variance induced by cell divisions. The heteroplasmy variance induced by cell divisions can even be controlled to sub-binomial levels in the case of self-avoidance, strong discrimination, and a homogeneous network distribution.

Notably, it is possible for the cell to control copy number variance below the binomial limit while also generating heteroplasmy variance, without biased network inclusion (for example, in the *n* = 64, *h* = 0.5 cases in Fig. 4), reflecting a potentially beneficial case for implementing a genetic bottleneck without challenging overall mtDNA levels.

Analytic progress is more challenging for this case, but an imperfect statistical model (Fig. 7; see Methods) can begin to capture some of the qualitative behaviour. The model correctly predicts the direction of change of copy number variance for various network structures, and the capacity to control variance below the binomial value, but the magnitudes of predicted variances are more extreme than those observed in simulation. This discrepancy arises because, to retain tractability, the algebraic model imposes an even spread of mtDNAs more strictly than is applied in the simulation (where this imposition is limited for numerical reasons). The range of variance values in the simulation is thus more limited than those that emerge from model predictions, although the trends of behaviour with governing variables are largely consistent.

### Diffusion of mtDNA relaxes statistics towards their null value

The mitochondrial network fragments before cell division. This fragmentation gives a time window during which mtDNAs that were previously constrained by the network structure can diffuse away from their initial position. In the limit of infinite diffusion, network structure will be forgotten and the mtDNA population will be randomly and uniformly distributed throughout the cell, leading to binomial inheritance patterns. We next investigated how limited amounts of diffusion away from the initial structure influence the patterns of mtDNA inheritance.

Fig. 5 shows *V′*(*h*) for random placement of mtDNA in the network (left column) and repulsive placement of mtDNA in the network (right column) for 4, 16, and 64 seed points (more seed points give a more homogeneous network), respectively, in rows from the top to the bottom. Diffusion is applied through 100 normally-distributed random steps of width λ, so that mtDNA molecules undergo random walks with expected total displacement around 10λ (though not exactly this value, due to boundary conditions). This model (Fig. 1G-H) mirrors findings that mitochondria may undergo a series of directed bursts of motion from active randomization prior to cell division [50]. Our results indicate that directed bursts of motion of the mitochondria indeed work to remove the effects of the network on daughter cell statistics, resulting in binomial segregation. Notably, there are cases in which extremely heterogeneous mitochondrial networks can leave an imprint on mtDNA inheritance despite high diffusion strength (top row).

## Discussion

We have demonstrated that a cell’s mitochondrial network structure can control cell-division variability in both mtDNA copy number and heteroplasmy inheritance, in both directions (Fig. 6). When different mtDNA genotypes have different propensities for network inclusion, random arrangement in a heterogeneous network generates (much) more variability than could be achieved through random cytoplasmic arrangement alone. On the other hand, ordered arrangement in a homogeneous network can control heteroplasmy variance below the binomial level expected from random partitioning. In concert, homogeneous network structure dramatically reduces copy number variance to sub-binomial levels (as observed experimentally in Ref. [53]), and heterogeneous network structure correspondingly increases it. Notably, it is possible to tightly control copy number while generating heteroplasmy variance – a strategy that may be useful in implementing a beneficial genetic bottleneck to segregate mtDNA damage.

How do mitochondria in different taxa and tissues correspond to the different regimes in our model? In animals, mitochondrial structure is highly varied, from largely fragmented organelles (low *p* and *q* in our model) to highly reticulated networks (high *p* and/or *q*), with mtDNA nucleoids moving freely throughout the mitochondrial reticulum, leading to a random distribution of mtDNA within the network [57, 58]. Animal mitochondria are expected to fragment prior to cell divisions, with a spectrum of active randomization mechanisms [59, 50], captured by the diffusion behaviour in our model. Fungal mitochondria, on the other hand, are often inherited without fragmenting the network, which remains in its reticulated state through cell division [32]. *Saccharomyces cerevisiae* populations have been shown to clear heteroplasmy within a few generations, with relatively heterogeneous networks and semi-regular spacing of mtDNA [24]. This situation corresponds to the ‘repulsive’ version of our model, inducing heteroplasmy variance (helping to clear heteroplasmy) while controlling copy number. In plants, mitochondria normally exist in a fragmented state (low *p* and *q*) [60, 61]. An intriguing exception to this is the formation of a reticulated network prior to division in the shoot apical meristem – the tissue that gives rise to the aboveground germline [18]. This formation could be the signature of network structure being employed to shape mtDNA prior to inheritance – although this employment is also likely to involve facilitating recombination [16].

One technical observation from our work is that the Taylor expansion method for approximating heteroplasmy variance, often employed in mtDNA models [27, 55], has several shortcomings in the face of network structure and other physical phenomenology that induce correlations between mtDNA types. Higher-order terms in the expansion do not immediately fix the discrepancies from simulation; we conclude that the series is likely slow to converge in these cases. Our model does successfully capture all the moments and covariances of the quantities of interest (see Appendix) – so, for example, all pertinent statistics of the number of wildtype mtDNAs can reliably be extract. It is the combination of these statistics into an approximation for a ratio (heteroplasmy) that is unreliable. We advocate the use of analytic expressions (like our exhaustive sum over states) and explicit simulation to check the validity of such results in future contexts (in a sense paralleling the careful consideration of moment-based methods for stochastic chemical kinetics [62]).

Our observations on how network structure influences genetic population structure stand in parallel with the many other phenomena associated with the physical and genetic behaviour of mitochondria. Mitochondrial network structure and dynamics likely fulfil many purposes [34], including contributing to mtDNA quality control [44, 43] via facilitating selection. Here, we assume that selection occurs (if at all) between cell divisions, focussing rather on the behaviour at cell divisions. Previous modelling work has demonstrated the capacity of mitochondrial network structure to shape mtDNA genetics through ongoing processes through the cell cycle [49, 16]; other work has considered the behaviour of controlled mtDNA populations across divisions without considering how that control may be physically manifest [52, 56]. We hope that our models here help bridge the gap between these pictures of mitochondrial spatial dynamics between, and well-mixed behaviour at, cell divisions across a range of eukaryotic life.

## Methods

### Network simulation

We constructed a random network via an elongation and branching process; network segments elongated deterministically with rate *e* = 0.01 and branched according to Poissonian dynamics with a given rate *k* = 0.02, and terminated if they hit the cell boundary. Network growth was initiated at a number of evenly-spaced seed points around the cell circumference (the first of which was randomly positioned in each simulation). Initial segments then grew perpendicular to the perimeter of a circular 2D cell, represented by the unit disc. Network growth proceeded until a predetermined network mass had been created; if all segments terminated before this mass was reached, we re-seeded the perimeter, and continued the growth process. By changing the number of seed points from which segments grow, we tune the uniformity of the network structure: a high number of seed points yielded homogeneous network structures, whereas a low number of seed points often lead to heterogeneous network structures (see Fig. 1).

Next, we distributed mtDNAs in the cell. We considered two mtDNA types, wildtype *W* and mutant type *M*, each a predetermined proportion of the mtDNA population, as specified by *h*. Proportions *p* of wild type mtDNA and *q* of mutant type mtDNA molecules were then placed within the network according to a particular placement rule (see below). The remaining mtDNA molecules were randomly and uniformly distributed in the cell, modelling presence of fragmented organelles in the cytoplasm. First, we considered random and uniform placement of mtDNA within the network, in which every point of the network was equally like to host an mtDNA molecule. Later we introduce a minimal inter-mtDNA distance within the network, hence enforcing spacing between mtDNA molecules through a mutual repulsion.

Finally, we partition the cell and record the number of wildtype *W* and mutant *M* mtDNAs in one daughter, the heteroplasmy *h* = *m*/(*w* + *m*), as well as the proportion of network mass *u*. The process of cell division was modelled by recording only the network mass and mtDNA content of a fixed circular segment spanning an angle *ϕ*, randomly orientated with respect to the network seed points. We are particularly interested in the heteroplasmy variance *V*(*h*) and copy number variance *V*(*N*), where *N* = *W* + *M*, across many realisations of this system.

### Statistical models of mtDNA copy number and heteroplasmy

We consider 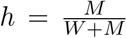 and *N* = *W* + *M* as our key variables. We assume that the parent cell’s heteroplasmy level is *h* ∈ [0, 1], with a total of *N*_0_ mtDNA molecules. Thus there are *hN*_0_ mutant molecules, and (1 – *h*)*N*_0_ wild type molecules of mtDNA.

We ignore correlations between daughter cells and focus on a single daughter from a cell division. In the daughter, the copy number variance is

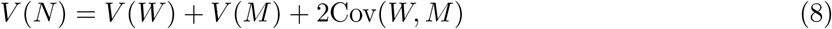

Here *V*(*W*) and *V*(*M*) are the variances of wild-type and mutant mtDNA, respectively, and Cov(*W*, *M*) is the covariance of *W* with *M*. The heteroplasmy variance, *V*(*h*) does not follow a simple form as it deals with a ratio of random variables. Instead, we use either explicit sums for the moments as in the text:

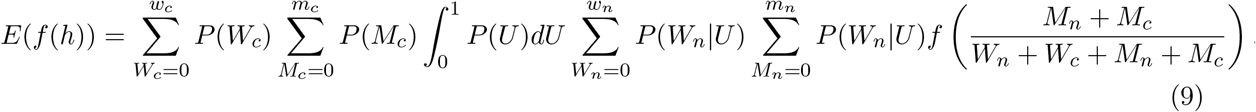

with mean *E*(*h*) and variance *V*(*h*) = *E*(*h*^2^) – *E*(*h*)^2^, or a first-order Taylor expansion, finding that

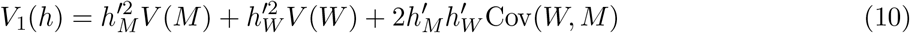

The prefactors 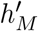 and 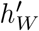 are derivatives of the heteroplasmy level considered as a function of *W* and *M*, and are model dependent (described below). Dividing *V*_1_(*h*) by *h*(1 – *h*), we get the normalized heteroplasmy variance, 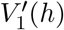, which conveniently removes some of the dependence on *h*. We also considered higher order terms in this expansion (see Appendix B.2).

### The null hypothesis: no mtDNA placement in network

As our null hypothesis, we considered a binomial segregation model for mtDNA [15]. In this model, no network structure exists and no active mechanisms contribute to the distributions of mtDNA (of either type) to the daughter cell. Supposing that the cell divides such that the daughter consists of a proportion *p_c_* of the parent cell volume, we supposed

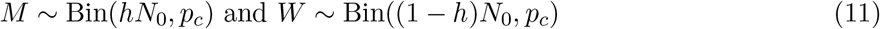

The copy number variance of the daughter is then, from the binomial distribution,

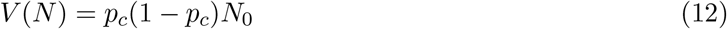

To find the heteroplasmy level variance, we calculated the variances of *W* and *M*, and the corresponding derivatives in Eqs. (24, 25); the covariance of *W* and *M* is zero in this case. A detailed derivation of *V*_1_(*h*) is presented in Appendix B.1, the result of which were

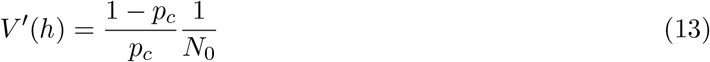

This is our null case, with nothing actively influencing the placement of mtDNAs within the cell. If this is the case, apportioning of mtDNA to daughter cells is binomial. The familiar expression of 1/*N*_0_ is recapitulated for symmetric cell divisions, i.e., with *p_c_* = 1/2, in which case *V′*(*h*) = 1/*N*_0_.

### Random mtDNA placement in network

Following intuition and preliminary observation of our simulations, we model the proportion *u* of network mass inherited by the smaller daughter as beta distributed variable *U*, with mean *E*(*U*) and variance *V*(*U*). Expected network inheritance *E*(*U*) will simply be *p_c_*, the proportion of cell volume inherited; *V*(*U*) will depend on the spread of the network through the cell, and constitutes a fit parameter in comparing this statistical model to simulation. Hence *u* is drawn from the beta distribution, Beta(*α*, *β*) with mean *E*(*U*) = *α*/(*α* + *β*) = *p_c_* and variance 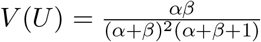.

We now write *W_n_*, *W_c_* respectively for the number of wildtype mtDNAs placed in the network and randomly spread in the cytoplasm, and *M_n_*, *M_c_* likewise for mutant mtDNA. *W_c_* and *M_c_* are assumed to follow the binomial partitioning dynamics above. Assuming that mtDNAs in the network are randomly positioned therein, we draw a *u* ~ Beta(*α*, *β*) to reflect the network proportion inherited by the smaller daughter, and write

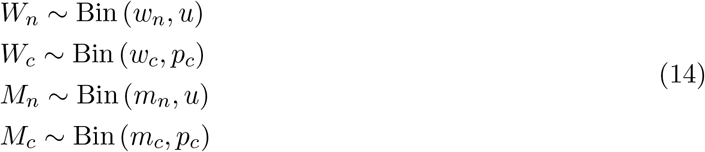

where *w_n_* = *p*(1 – *h*)*N*_0_, *w_c_* = (1 – *p*)(1 – *h*)*N*_0_, *m_n_* = *qhN*_0_, *m_c_* = (1 – *q*)*hN*_0_.

The mean and variance of *N* are readily derived using the laws of iterated expectation and total variance to account for the compound distribution of mtDNA in the network (Appendix). To estimate heteroplasmy variance, we combine the (co)variances of the different types with their respective prefactor (from Eqs. (26, 27) in Appendix B.1).

### Repulsive mtDNA placement in network

Next, we considered the case where mtDNAs placed in the network are not randomly positioned, but instead experience a repulsive interaction, and thus adopt a more even spacing. Capturing this picture perfectly with a statistical model is challenging; instead, we use the following picture. The proportion of inherited network mass *u* consists of a finite number of ‘spaces’, each of which can be occupied by at most one mtDNA molecule. Choose a number of spaces to fill, then sample mtDNA molecules from the available pool without replacement, assigning each drawn mtDNA to the next unoccupied network space. In this case, the final population of mtDNAs in the network is described by the hypergeometric distribution. We draw *u* ~ Beta(*α*,*β*) to reflect the network proportion inherited by the smaller daughter, and write

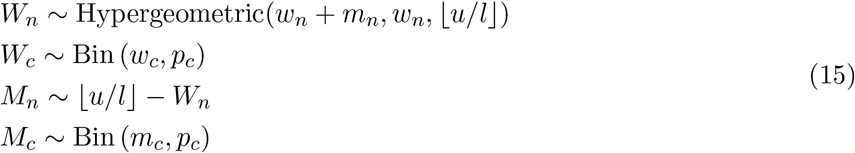

We can once again use the laws of total variance and iterated expectation (see Appendix B.1) to estimate heteroplasmy and copy number behaviour. As previously discussed, this model has several shortcomings and is only expected to match qualitative behaviour (see Appendix B.1).

## Code Availability

Code for this project is available at https://github.com/StochasticBiology/mtdna-network-partition

## Acknowledgments

This project has received funding from the European Research Council (ERC) under the European Union’s Horizon 2020 research and innovation programme (Grant agreement No. 805046 (EvoConBiO) to IGJ).

## Appendices

## A Models of mtDNA copy number variance

We consider four state variables describing the mtDNA population of a daughter cell after a mother divides: *W_n_*, *W_c_*, *M_n_*, *M_c_*, for the wildtype (*W*) and mutant (*M*) mtDNAs contained in a reticulated mitochondrial network (_*n*_) or in fragmented mitochondrial elements in the cytoplasm (_*c*_). An additional variable *U* describes the proportion of network mass inherited by the daughter cell.

The mother cell has *N*_0_ mtDNAs, a proportion *h* of which are mutants (*h* is heteroplasmy). Proportions *p* and *q* of the wildtype and mutant mtDNAs are contained in the network; the remainder are in fragments in the cytoplasm. We consider a daughter inheriting a proportion *p_c_* of the cytoplasm. The mtDNA copy number of the daughter, *N* = *W_n_* + *W_c_* + *M_n_* + *M_c_*, is the sum of all components. We write *W* = *W_n_* + *W_c_* and *M* = *M_n_* + *M_c_*. Under the null hypothesis of no network, since *M* ~ Bin(*hN*_0_,*p_c_*) and *W* ~ Bin((1 – *h*)*N*_0_,*p_c_*), we have *N* ~ Bin(*N*_0_,*p_c_*) and

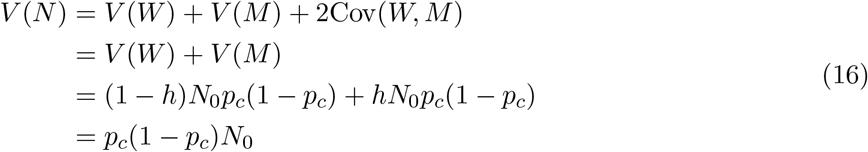

For the random mtDNA distribution model, using the full model (Eq. 1), and decomposing into the networked and individual mitochondria, i.e., *N* = *N_c_* + *N_n_*, the law of total variance gives

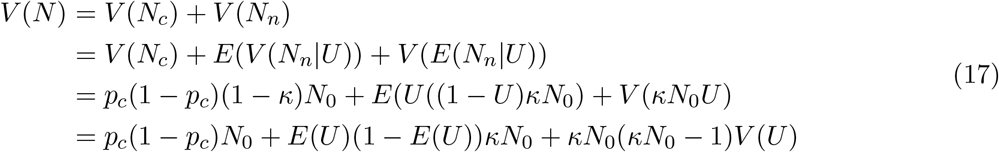

Overall, this expression captures copy number variance dynamics across a wide range of parameterisations (see Fig. 9). For the repulsive mtDNA distribution model (15) the corresponding expressions are

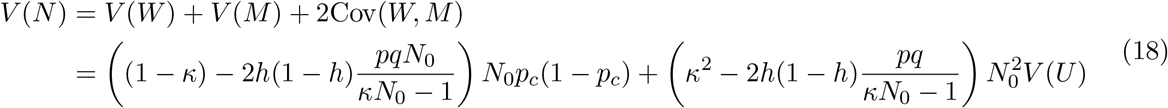

where *κ* = *p*(1 – *h*) + *qh*. The expression correctly predicts sub-binomial variance for heterogeneous networks, but fails to capture the structure of genetic bias as well as other qualitative dynamics of the system (see Fig. 10).

**Figure 9:**
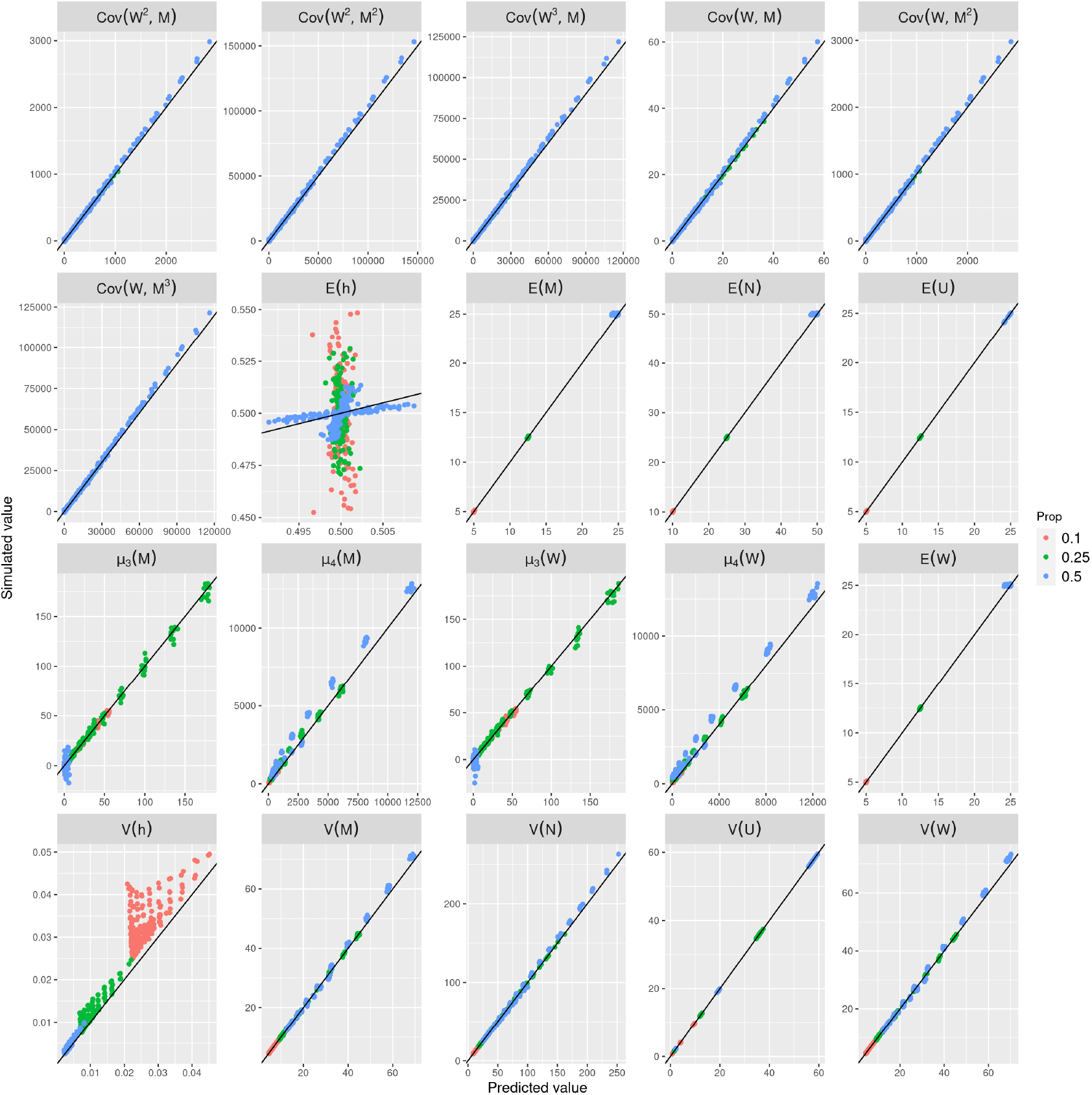
Comparisons of individual simulation moments (vertical axes) and analytic moments (horizontal axes) for random placement of mtDNA in the network (Eq. 1) for varying proportions of cell volume inherited by the smallest daughter. Colors reflect the proportion of parent cell volume apportioned to the daughter of interest.

**Figure 10:**
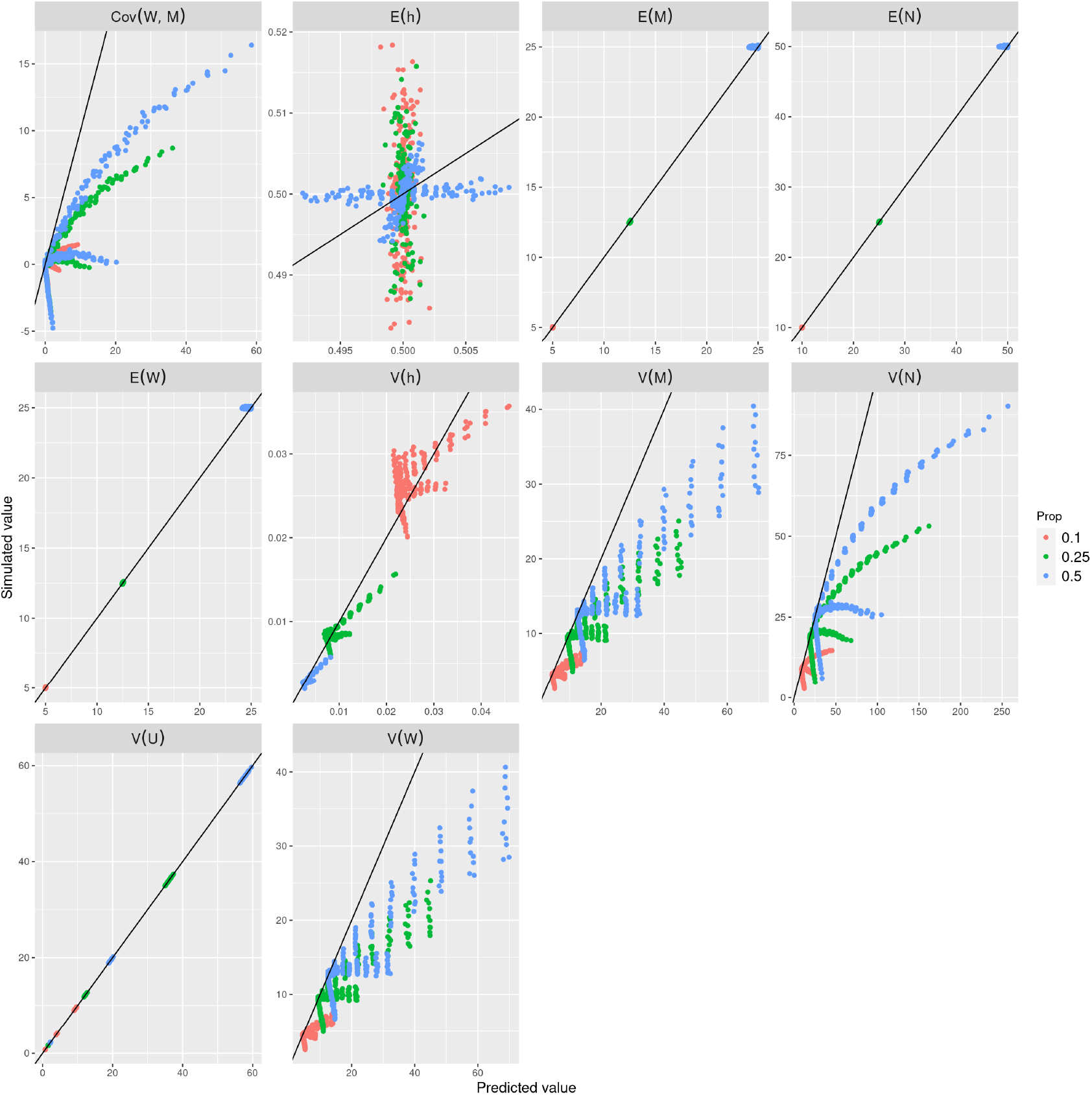
Comparisons of individual simulation moments (vertical axes) and analytic moments (horizontal axes) for repulsive placement of mtDNA in the network (Eq. 15) for varying parent cell cytoplasm proportions inherited by the smallest daughter. Colors reflect the proportion of parent cell volume apportioned to the daughter of interest.

## B Heteroplasmy level variance

The heteroplasmy level is the mutant proportion of the cell, i.e.,

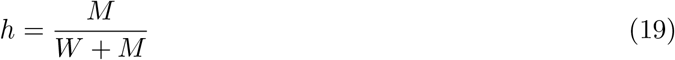

where *W* and *M* are the numbers of wild-type and mutant type mtDNA, respectively. By definition,

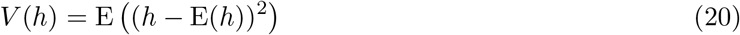

and *h* = *h*(*M*, *W*). As the ratio of random variables, *h* does not admit as straightforward an analysis as *N*. Using a sum over all states of the system (Eq. 2) in the main text we can compute its value (and *V*(*N*)) for a given system. While these expressions provide a good match with simulations for a wide range of parameterisations (see Fig. 3), they do not allow intuitive understanding of the system. Since we sought a more intuitive analysis, we employ first- and second-order Taylor expansions to generate more tractable estimations focussing on key governing parameters.

### B.1 Taylor expansions for heteroplasmy level variance

Using a first-order Taylor expansion (as in, for example, Ref. [27]), we obtain

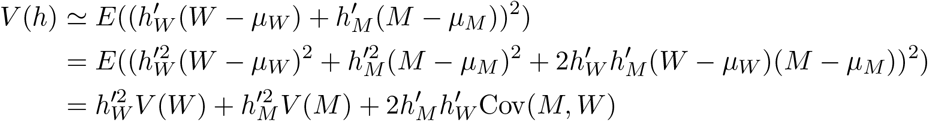

where prefactors 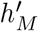 and 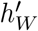 are the partial derivatives of *h*(*M*, *W*) evaluated at the means of the distribution for *M* and *W*, *μ_M_* = *hN*_0_ and *μ_W_* = (1 – *h*)*N*_0_. In general, we find

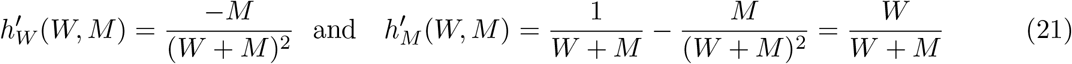

or

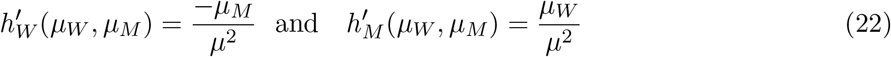

This approximation is used to derive the variance of the heteroplasmy level in all the scenarios considered, so we give it an subscript of 1 to show it is a first-order Taylor expansion.

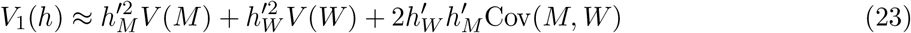

The evaluation of the prefactors are model-specific: Under the null hypothesis, where both *W* and *M* are binomially distributed with probability *p_c_* and their respective proportion of the population *N*, *μ_M_* = *p_c_hN*_0_, *μ_W_* = *p_c_*(1 – *h*)*N*_0_ and *μ* = *μ_M_* + *μ_W_*, these expressions evaluate to

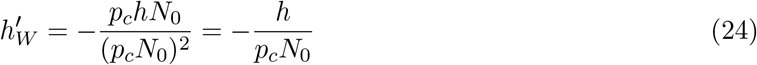

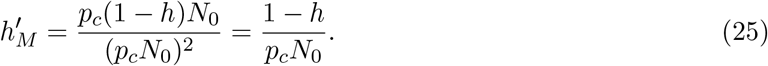

For the network model, whether it is with or without mutual repulsion of mtDNA molecules, we write *μ_W_* = *μ_W_n__* + *μ_W_c__* = *E*(*U*)*w_n_* + *p_c_w_c_* and *μ_M_* = *μ_M_n__* + *μ_M_c__* = *E*(*U*)*m_n_* + *p_c_m_c_*, so *μ* = *μ_W_* + *μ_M_* = *E*(*U*)(*w_n_* + *m_n_*) + *p_c_*(*w_c_* + *m_c_*) and

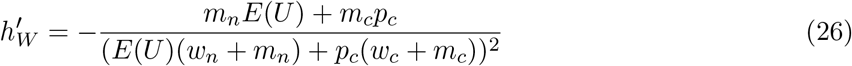

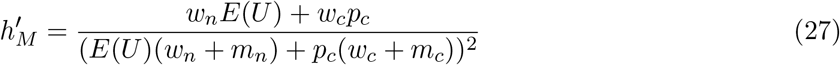

Note that under the assumption that the network is evenly distributed throughout the cell, i.e., *E*(*U*) = *p_c_*, we recapitulate the expressions for the null hypothesis pre-factors.

#### Null model (no network structure)

Under the null hypothesis, independent binomial distributions describe both *M* and *W*. Combining Eqs. 24, 25) with the variances of each specie, we find

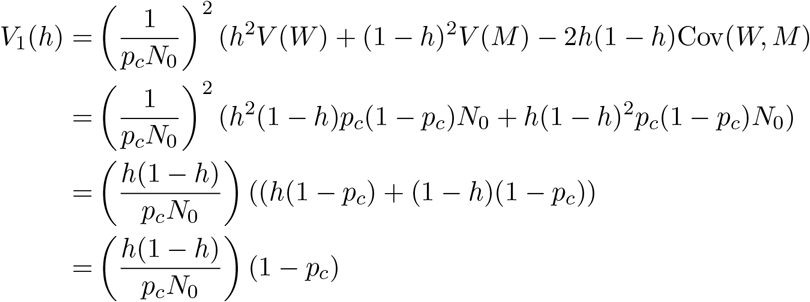

which, weighted by *h*(1 – *h*), gives Eq. 13, i.e., the normalized heteroplasmy variance defined as

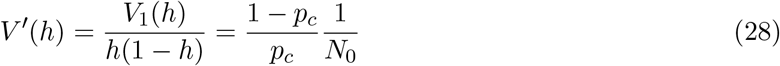

#### Network model without repulsion

We next consider the influence of network structure on mtDNA distributions. All random variables are binomially distributed with their respective proportion of the total mtDNA content of the parent as the population, with *p_c_* or *u* as the probability for cytoplasmic and networked mtDNAs respectively. Using the law of total variance for *M* and *W*, we find that

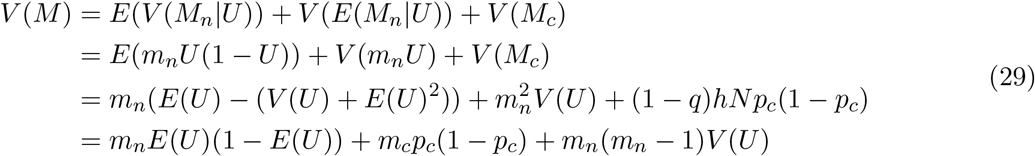

and

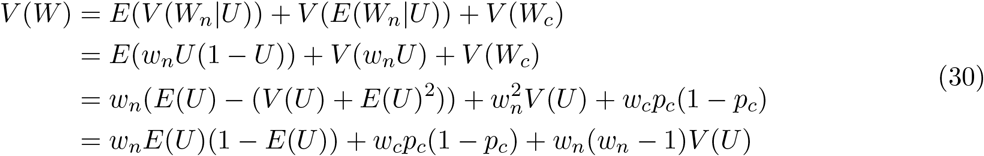

Using the law of total covariance, since cytoplasmic copy numbers are uncorrelated, we find that the covariance of *M* with *W* is

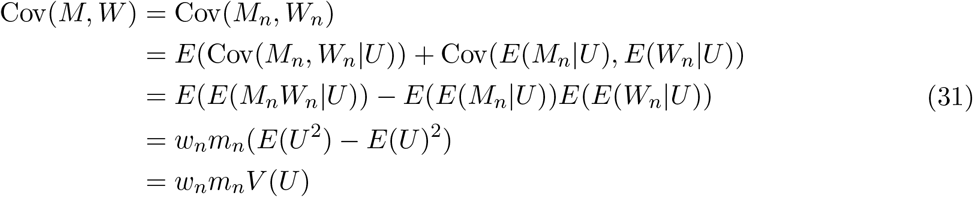

Combining (co)variances with the prefactors of Eqs. (27,26), Eq. (23) gives

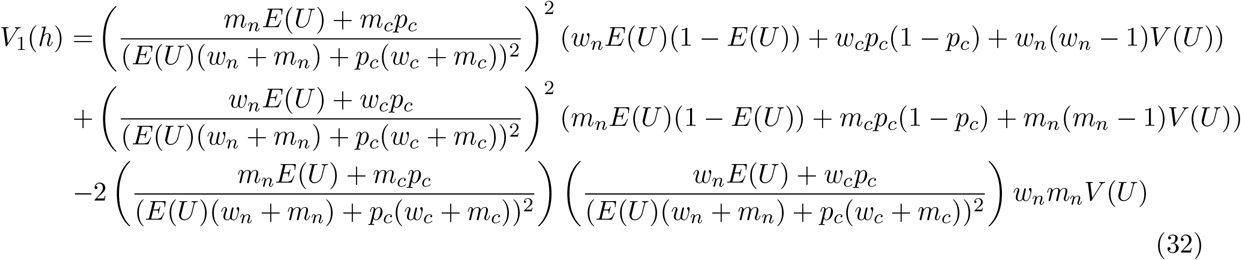

If we assume that *E*(*U*) = *p_c_*, for which *w_n_* + *w_c_* = (1 – *h*)*N*_0_ and *m_n_* + *m_c_* = *hN*_0_, we get a simpler expression,

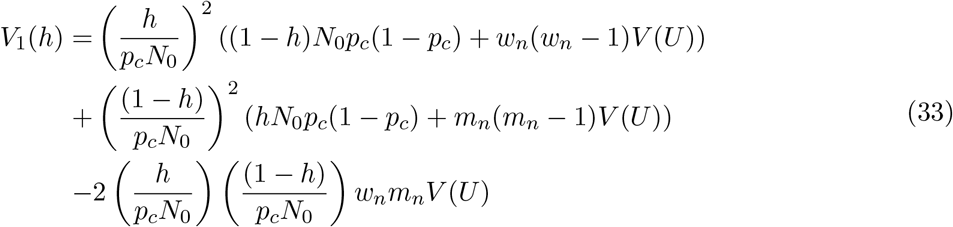

Simplifying and gathering terms, we find that

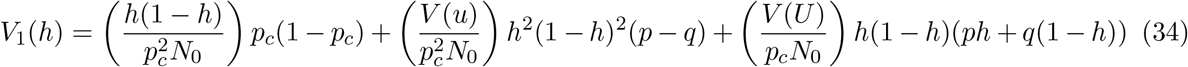

Dividing by *h*(1 – *h*) gives 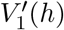,

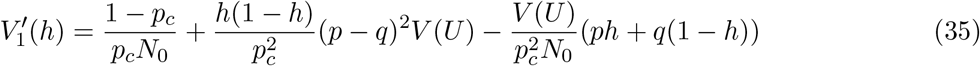

which we plot for different values of *p*, *q*, *p_c_*, and network parameters, *E*(*U*) and *V*(*U*) taken from simulation. The latter two are used to fit a beta distribution with parameters *α* and *β*, with which we model the process of network segregation when a cell divides.

Looking to gain insights from Eq. 35, we write *p* = *q* – *δ*. In this case, supposing that *δ* is small, wild-type and mutant mtDNA are treated almost equally by the network, with an almost equal proportion of the two admitted into the network.

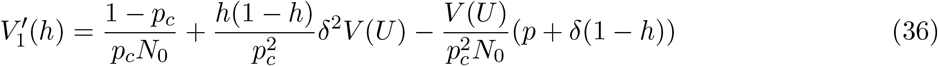

It will be seen that there is a quadratic dependence on *δ*, the difference in inclusion probabilities for the different types of mtDNA. For *δ* ≠ 0, the network is genetically biased towards one of the types, to which there is associated an increase in *V′*(*h*). For *δ* = 0, the network is genetically unbiased, giving

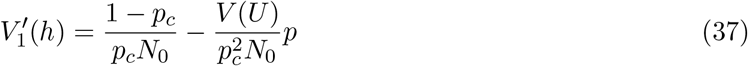

When *p_c_* = 1/2, i.e., when cell division is symmetric, we arrive at Eq. 4 in the main text. Eq. 37 suggests that the network structure provides a negative contribution to *V′*(*h*), resulting in the low diagonal values in the first order Taylor expansion of *V′*(*h*), whereas, from the simulations, we expected a small *increase* along the diagonal (shown in Fig. 2). The second order Taylor expansion corrects this (Fig. 8), with contributions of third order and higher in p and q (Eq. 46), but it overcompensates; we do not pursue higher order terms, mostly because they are hard to interpret, and present significant difficulties in calculations. We then asked whether statistical simulations would produce a better fit, and we find that there is good support for this model when mtDNA molecules are randomly distributed throughout the network in our simulations.

#### Network model with repulsion

Next we considered networks in which mtDNA molecules within the network were mutually repulsed by each other, setting a minimum distance between mtDNA molecules in the network. We again decompose *W* and *M* into their cytoplasmic and network components, i.e.,

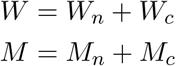

To assess the effect of mutual self-repulsion of mtDNA, we assumed a model of mtDNA transmission from a parent to its smaller daughter in which we consider the network to be divided into ⌊*u/l*⌋ different compartments. Into each of these compartments a single mtDNA molecule will be placed – hence *l* acts to enforce a minimum inter-mtDNA distance due to the repulsion of mtDNA molecules. We then fill these places by sampling, without replacement, mtDNA molecules from the set contained in the network. This corresponds to a hypergeometric model of ⌊*u/l*⌋ samples from a population of *w_n_* + *m_n_* mtDNAs, *w_n_* of which are wild-type,

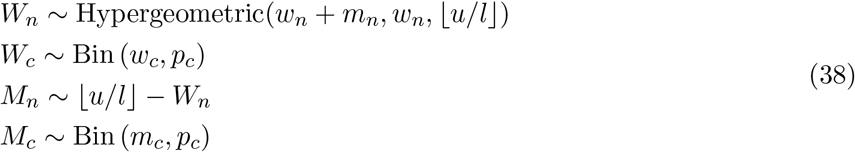

where *u* ~ beta(*α*, *β*). The problem with this model is that, to keep the values of *W_n_* and *M_n_* consistent, we must assume that there are exactly *w_n_* + *m_n_* spaces in the network; hence *l* = (*w_n_* + *m_n_*)^-1^. In our physical simulations, l is instead set to a distance (*l* = 0.01) that enforces some separation between mtDNAs while making it possible to populate the network through random positions in reasonable time. There are therefore more ‘spaces’ in the simulation than captured by the model, meaning that an even physical spread is less enforced in the simulation than in the model, and the range of variance values supported will be more limited in the simulation.

Accepting this limitation, the variance of *W_n_* is derived using the law of total variance, giving

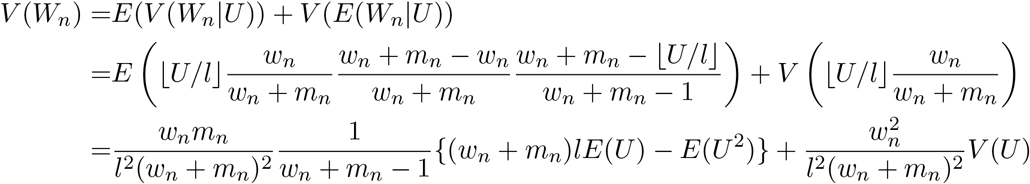

And using the *l* = (*w_n_* + *m_n_*)^-1^ assumption,

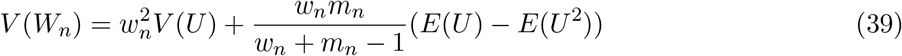

For *V*(*M_n_*) we find

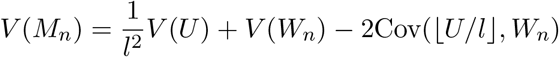

Using the law of total covariance,

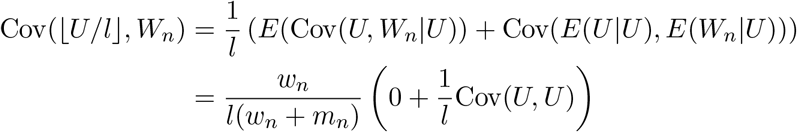

Setting *l* = (*w_n_* + *m_n_*)^-1^, we find

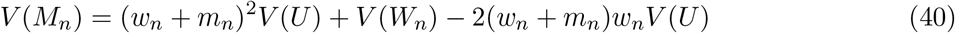

Combining with the variances of the cytoplasmic mtDNA content, we find that

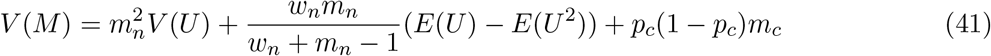

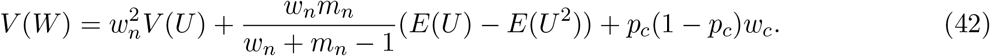

As before, *W_n_* has non-zero covariance with *M_n_*, due to their mutual dependence on network structure, but no cytoplasmic component covaries with any other component. The overall mutant-wildtype covariance is therefore

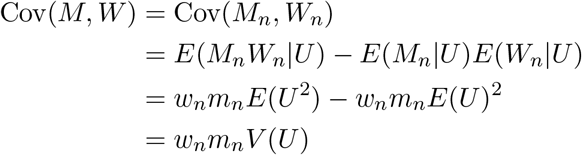

In this case, *V*_1_(*h*), the first-order Taylor expansion of heteroplasmy variance is

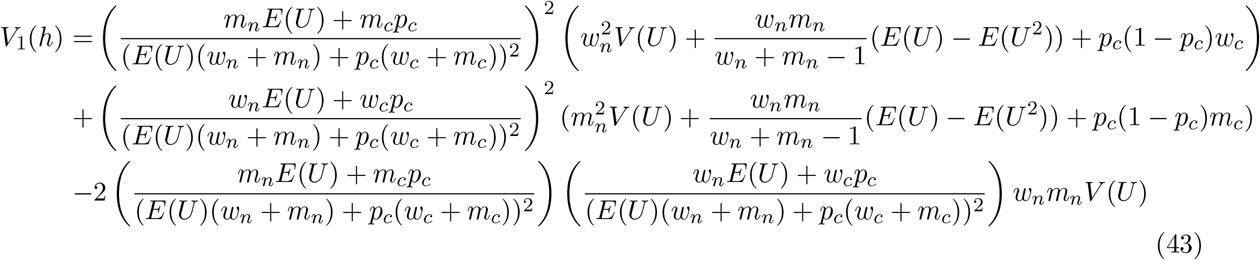

Assuming *E*(*u*) = *p_c_*, we find that

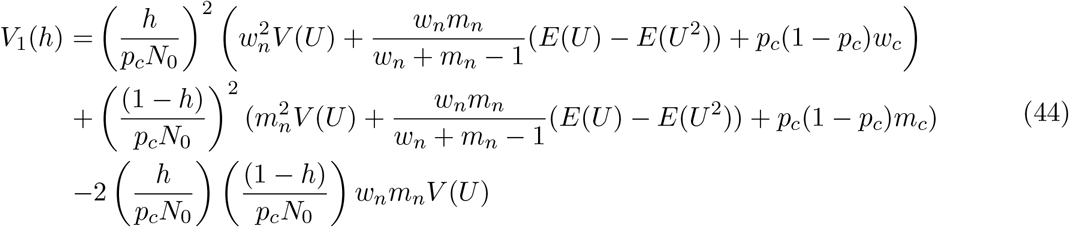

Gathering some terms and dividing by *h*(1 – *h*), we find

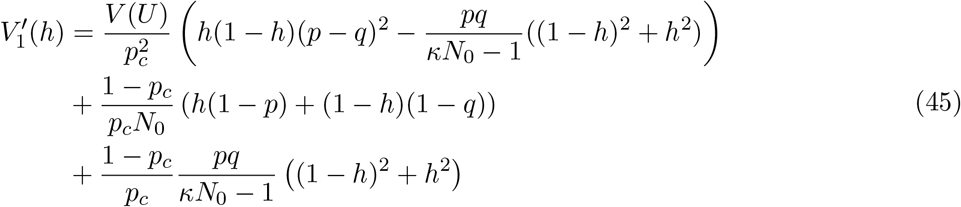

where *κ* = *p*(1 – *h*) + *qh*. As before, we plot 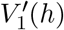 for different values of *p*, *q*, *p_c_*, and given network parameters *E*(*U*) and *V*(*U*), used to fit a beta distribution with parameters *α* and *β*, with which we model the process of mtDNA distribution when a cell divides. Fig. 7 shows the result of plotting 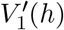 for networked distributions with mutually repulsive mtDNA molecules. Despite imperfections (Fig. 10), it will be seen that the the model captures qualitative behaviour of the simulations.

### B.2 Higher-order moments and second-order Taylor expansion

Given some observed shortcomings in the ability of the first-order Taylor expansion to capture heteroplasmy variance, we asked whether the next-order terms in the Taylor expansion could refine the estimates. The second-order Taylor expansion of heteroplasmy level variance used for the nonuniform distribution mtDNAs can be expressed as *V*_1_(*h*) + *V*_2_(*h*):

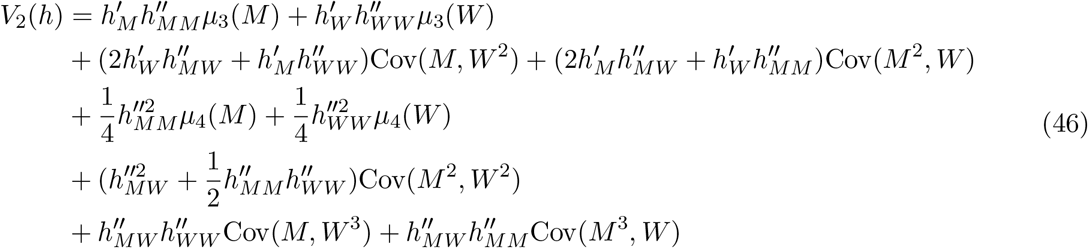

where *V_1_*(*h*) is given by Eq. 4, and the derivatives are given by

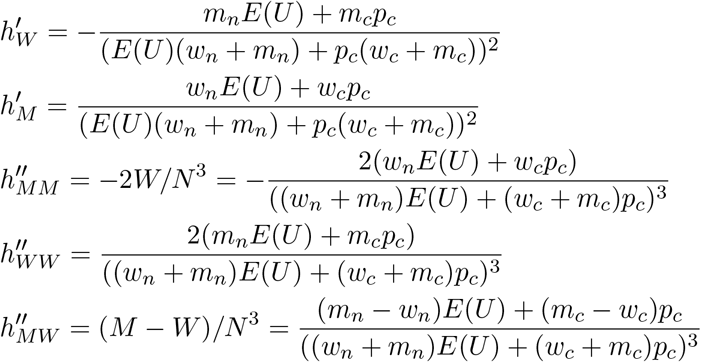

#### B.2.1 Higher-order moments from binomial and beta-binomial distributions

Expanding the Taylor expansion to second-order, we need a number of higher-order moments of the distributions of *W* and *M*. We start by calculating the third and fourth central moments of *W* and *M*. For the third order moments, we write

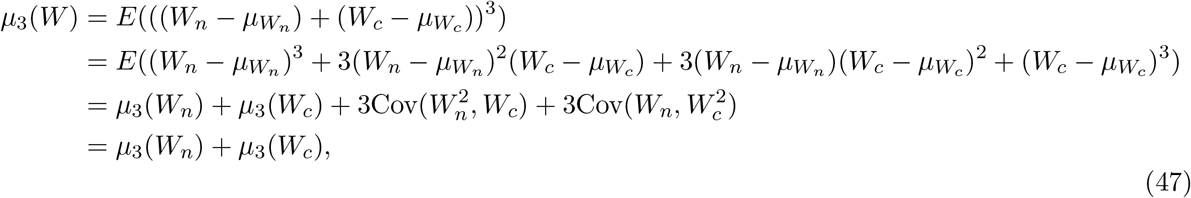

where the final line follows because networked and cytoplasmic mtDNA counts are uncorrelated. As *W_n_* is beta-binomial, we can take established expressions for the moments of the beta-binomial distribution; for *W_c_* we use established expressions for the binomial distribution [63]:

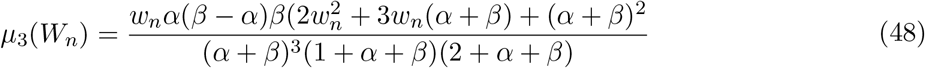

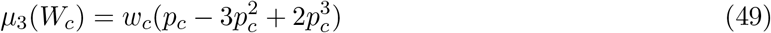

The fourth central moment of *W*n** is taken from the beta-binomial distribution:

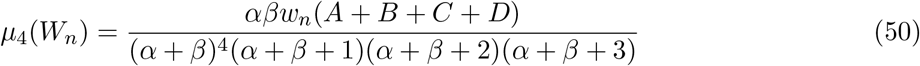

where

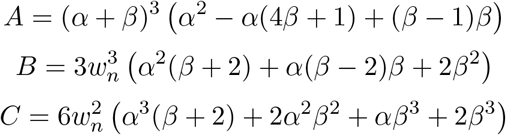

and

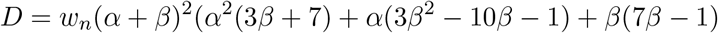

The fourth central moment of *W_c_* is from the binomial distribution:

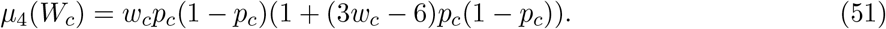

For *M* we find the same expression, but with different prefactors

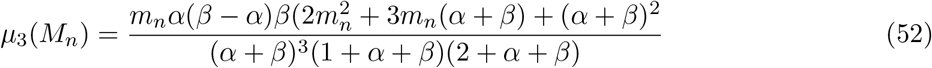

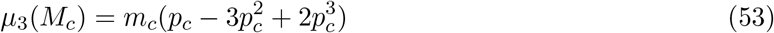

For terms in *M* we follow the same approach of recruiting central moment results from the betabinomial and binomial distributions. The same expression structures arise, but with different prefactors reflecting the mutant population:

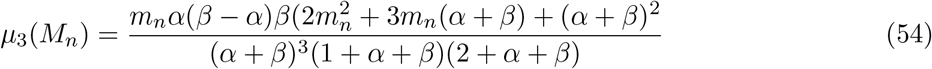

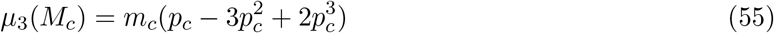

and

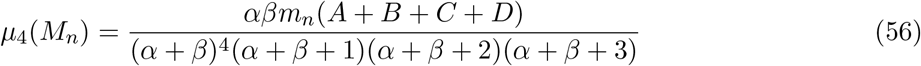

where

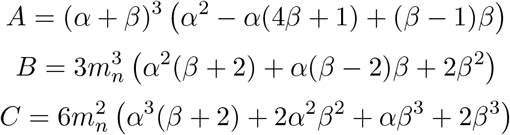

and

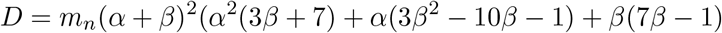

The fourth central moment of *M_c_* is

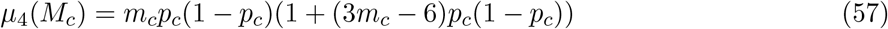

#### B.2.2 Covariance calculations

To calculate the covariances, we use the law of total covariance, which for RVs *X*, *Y* and *Z* states that

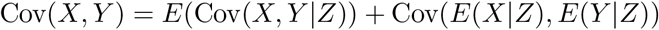

Using the identity Cov(*X*, *Y*) = *E*(*XY*) – *E*(*X*)*E*(*Y*), we find we retain the terms

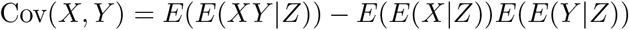

In this case, when *u* is fixed, the variables *W* and *M* are independent RVs, so the first term is the *E*(*E*(*X*|*Z*)*E*(*Y*|*Z*)). Using these findings, the necessary covariances of higher order in the RVs *M* and *W* are

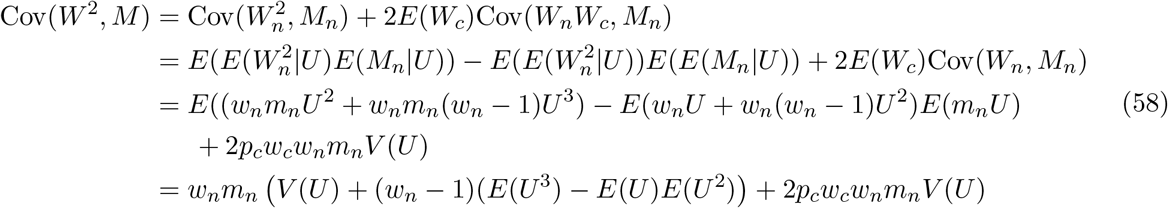

Note that we have used that 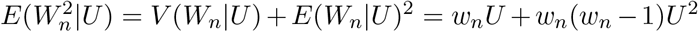. To calculate the covariance of *W*^2^ with *M*^2^, we also need 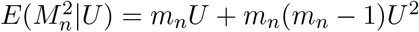

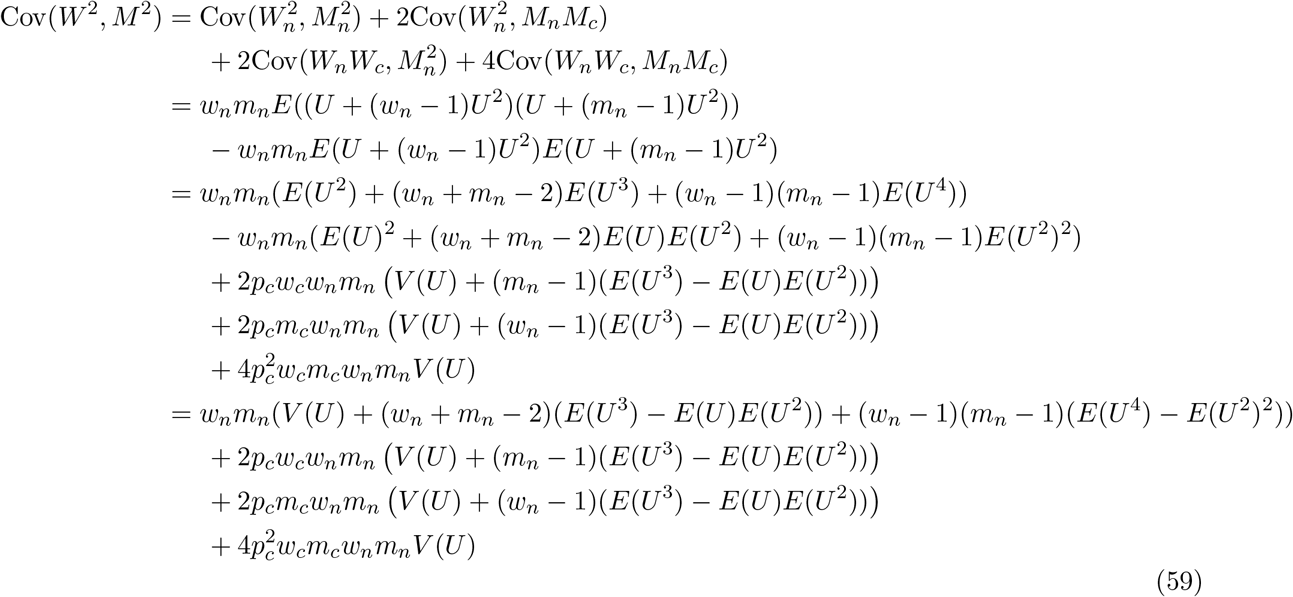

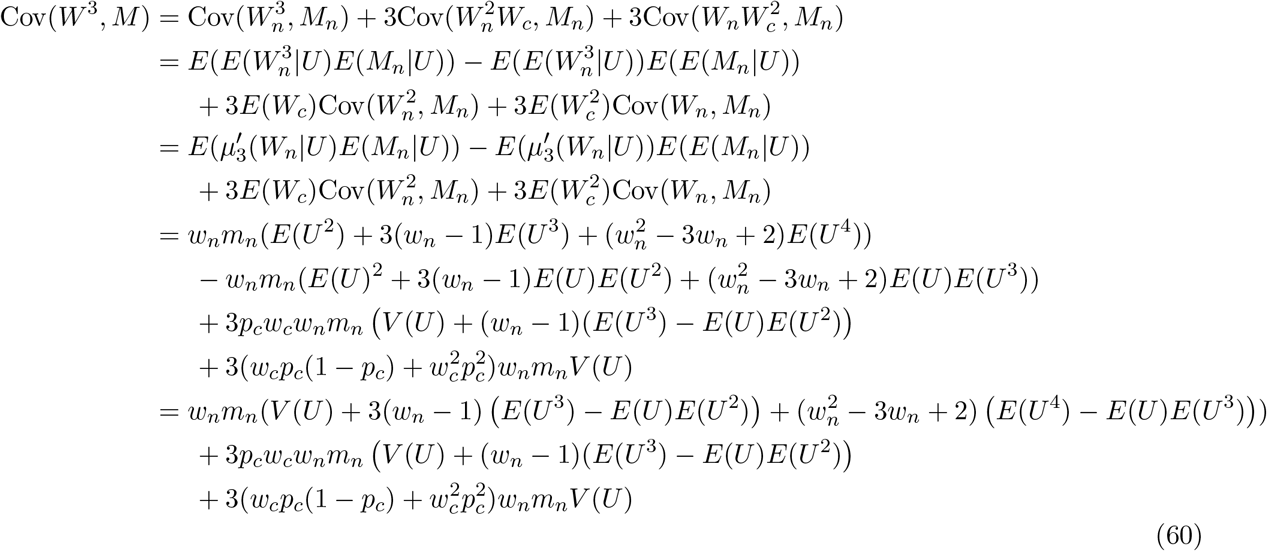

Here we have used 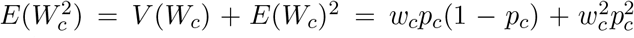 and, since the third non-central moment 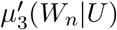 expressed in terms of the third central moment *μ*_3_(*W_n_*|*U*) is 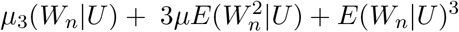, where *μ*_3_(*W_n_*|*U*) = *w_n_*(*U* – 3*U*^2^ + 2*U*^3^), we find that

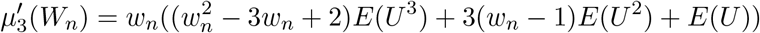

Lastly, Cov(*W*, *M*^2^) and by the symmetry of the problem

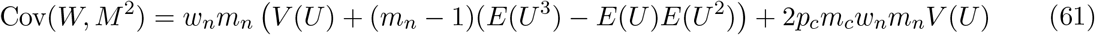

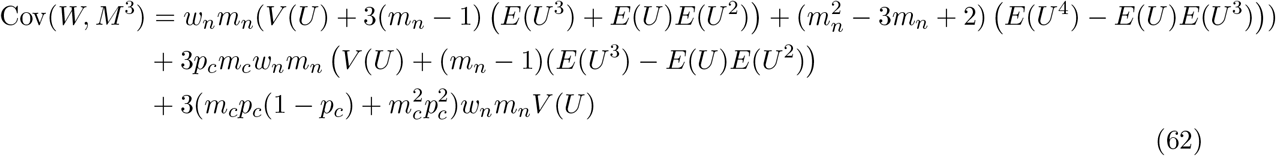

In Figs. 9–10, we plot these expressions for the various moments and covariances in the system compared to those arising from our simulation model. We generally observe good agreement between theory and simulation (most departures are on a very small scale compared to the overall scale of the corresponding expression; arising due to small deviations from the beta-distribution model for network mass). However, the overall second-order Taylor expression still deviates substantially from observed heteroplasmy variance (Fig. 8). One aspect of the first-order model is improved – the increase on the *p* = *q* diagonal. But the off-diagonal behaviour is substantially compromised, suggesting an overcompensation to the errors in the previous order. We conclude that higher-still terms in the expansion will be required to more perfectly capture the behaviour, and that convergence to the true behaviour may be rather slow.

#### B.2.3 Comparison of individual statistics

Here, we present the comparisons of simulation results with model results in both models to all relevant orders. First, one should note that the second order result only applies to the models with random mtDNA placement in the network, and then only for the heteroplasmy variance. This is because *h* is a ratio of random variables, and which is differentiable an arbitrary number of times with respect to both variables, *W* and *M*; the copy number variance, however, is linear in these random variables, and so the approximation is the same for all orders starting at first. Fig. 8 shows comparisons of simulation results (top row) with first and second order results (middle and bottom rows), respectively. Here we see clearly that both first and second order approximations were needed to capture the behavior displayed in our simulations, but that neither provides a reasonable match: the first order approximation departs significantly along the diagonal, displaying a small decrease as opposed to a small increase; the second order approximation massively overestimates, causing a far too large an increase along the diagonal.

